# Proteomic analysis of metabolic pathways supports chloroplast-mitochondria cross-talk in a Cu-limited diatom

**DOI:** 10.1101/2020.09.15.298752

**Authors:** Anna A. Hippmann, Nina Schuback, Kyung-Mee Moon, John P. McCrow, Andrew E. Allen, Leonard F. Foster, Beverley R. Green, Maria T. Maldonado

**Author notes:** **Author for Contact details:**, Dr. Anna A. Hippmann, University of British Columbia, Department of Earth Ocean and Atmospheric Science, Earth Sciences Building, 2207 Main Mall, Vancouver, V6T 1Z3, Canada.

## Abstract

Diatoms are one of the most successful phytoplankton groups in our oceans, being responsible for over 20% of the Earth’s photosynthetic productivity. Their chimeric genomes have genes derived from red algae, green algae, bacteria and heterotrophs, resulting in multiple isoenzymes targeted to different cellular compartments with the potential for differential regulation under nutrient limitation. The resulting interactions between metabolic pathways are not yet fully understood.

We previously showed how acclimation to Cu limitation enhanced susceptibility to overreduction of the photosynthetic electron transport chain and its reorganization to favor photoprotection over light-harvesting in the oceanic diatom *Thalassiosira oceanica* (Hippmann et al., 2017). In order to gain a better understanding of the overall metabolic changes that help alleviate the stress of Cu limitation, we have further analyzed the comprehensive proteomic datasets generated in that study to identify differentially expressed proteins involved in carbon, nitrogen and oxidative stress-related metabolic pathways.

Metabolic pathway analysis showed integrated responses to Cu limitation. The up-regulation of ferredoxin (Fdx) was correlated with up-regulation of plastidial Fdx-dependent isoenzymes involved in nitrogen assimilation as well as enzymes involved in glutathione synthesis, thus integrating nitrogen uptake and metabolism with photosynthesis and oxidative stress resistance. The differential expression of glycolytic isoenzymes located in the chloroplast and mitochondria may enable them to channel both excess electrons and/or ATP between these compartments. Additional evidence for chloroplast-mitochondrial cross-talk is the increased expression of chloroplast and mitochondrial proteins involved in the proposed malate shunt under Cu limitation.

**One sentence summary:** Diatoms adapt to Cu limitation by regulating their large repertoire of isoenzymes to channel electrons away from the chloroplast, enhance nitrogen uptake, and integrate the oxidative stress response. ^1,2,3^

## Introduction

Diatoms form an integral part of our oceans, influencing nutrient cycling and productivity of many marine foodwebs (Armbrust, 2009). Annually, marine diatoms fix as much carbon dioxide through photosynthesis as all terrestrial rainforests combined (Field et al., 1998; Nelson et al., 1995), thus having a significant impact on atmospheric CO2 levels and global climate. One key to their success may lie in their complex evolutionary history (Moustafa et al., 2009; Oborník and Green, 2005) which resulted in a mosaic genome with genes derived from the original heterotrophic eukaryotic host cell, the engulfed green and red algal endosymbionts, and a variety of associated bacteria (Armbrust et al., 2004; Bowler et al., 2008; Finazzi et al., 2010). As a result, diatoms possess multiple isoenzymes in many metabolic pathways, especially in carbon metabolism (Ewe et al., 2018; Gruber et al., 2009; Gruber and Kroth, 2014; Kroth et al., 2008; Smith et al., 2012).

The presence of multiple isoenzymes with different evolutionary histories also led to novel locations and interactions among metabolic pathways compared to green algal and animal ancestors (Allen et al., 2011; Gruber and Kroth, 2017). For example, in animals the complete set of proteins involved in glycolysis is located in the cytosol, whereas in green algae the first half of glycolysis (glucose to glyceraldehyde-3-phosphate, GAP) is located in the chloroplast and the second half (GAP to pyruvate) in the cytosol. In diatoms, an almost complete set of glycolytic proteins is found in both the cytosol and the chloroplast, with an additional set of proteins from the second half of glycolysis located in the mitochondria (Kroth et al., 2008; Río Bártulos et al., 2018; Smith et al., 2012). Furthermore, proteins involved in the ancient Entner-Dourodoff pathway, which is predominantly restricted to prokaryotes and catabolizes glucose to pyruvate, have also been identified in diatom genomes and are targeted to the mitochondria (Fabris et al., 2012; Río Bártulos et al., 2018).

A study by Allen et al (2012) illustrates the complexity of isoenzymes in diatoms further: the genome of *Phaeodactilum tricornutum* (*P. tricornutum*) encodes five different fructose-bisphosphate aldolase (FBA) isoenzymes, three targeted to the chloroplast and two to the cytosol (Allen et al., 2012). Each FBA has its own phylogenetic history. The expression pattern of these five isoenzymes changes depending on the nutritional status of the cell (Allen et al., 2012).

One of the most surprising discoveries from diatom genome sequencing was a complete urea cycle (Allen et al., 2011; Armbrust et al., 2004). In contrast to the catabolic nature of the urea cycle in animals, in diatoms it is an integral part of cellular metabolism and a hub of nitrogen and carbon redistribution within the cell. It is involved in amino acid synthesis, cell wall formation, carbon and nitrogen recycling, and it interacts with the citric acid cycle (Allen et al., 2011; Armbrust et al., 2004).

Most molecular studies on acclimation to nutrient limitation have focused on macronutrients, or on the essential micronutrient Fe, which limits phytoplankton in over 30% of the ocean (Moore et al., 2004). Some studies have shown an intricate interaction between Fe and Cu nutrition in phytoplankton (Annett et al., 2008; Guo et al., 2012; Maldonado et al., 2006, 2002; Peers and Price, 2006), but there are only a handful of studies on physiological adaptations to Cu limitation alone (Guo et al., 2015, 2012; Kong and M. Price, 2020; Lelong et al., 2013; Lombardi and Maldonado, 2011; Maldonado et al., 2006; Peers et al., 2005; Peers and Price, 2006).

Our recent comprehensive investigation on the physiological and proteomic changes to the photosynthetic apparatus of two strains of the open ocean diatom *Thalassiosira oceanica* (*T. oceanica*) in response to chronic Cu limitation revealed both similar and different strategies compared to those observed in response to low Fe (Hippmann et al., 2017). Acclimation to low Cu caused a bottleneck in the photosynthetic electron transport chain that was accompanied by major increases in the electron acceptors ferredoxin and ferredoxin:NADP+ reductase, which have major roles in counteracting reactive oxygen species. Along with changes in the composition of the light-harvesting apparatus, this resulted in a shift from photochemistry to photoprotection.

In our previous paper (Hippmann et al., 2017) we focussed on the photosynthetic electron transport chain and light-harvesting antennas as well as a number of physiological parameters changed in response to Cu limitation, but did not ask how carbon and nitrogen metabolism are affected and may interact when Cu is limiting. We now expand our proteomics analysis to include proteins involved in various carbon and nitrogen metabolic pathways (e.g. Calvin-Benson-Bassham cycle, glycolysis, TCA cycle, nitrogen acquisition and assimilation, urea cycle, malate shunt, glutathione metabolism), taking into account their predicted cellular compartments. Although the decrease in Rubisco activase suggests the CBB is down-regulated, there appear to be complex effects on the three-compartment glycolysis machinery. Increased expression of enzymes involved in nitrogen acquisition and assimilation could act simultaneously as a sink for reducing equivalents and as a supplier of compounds needed to support dissipation of reactive oxygen species. Finally, we present further evidence for cross-talk between chloroplast and mitochondria in form of an active malate shunt.

## Results

### Overview of proteomic datasets

In our original study (Hippmann et al., 2017), we investigated two strains, CCMP 1003 and CCMP 1005, of the centric diatom *T. oceanica* (here referred to as TO03 and TO05, respectively). On further examining the proteomic datatsets it was clear that Cu limitation had a stronger and more comprehensive effect on proteins of carbon and nitrogen metabolism in TO03 than T005 (this study), in line with observations for photosynthetic electron transport proteins Although the proteomic dataset of TO05 contains twice as many distinct proteins as that of T003 (1,431 versus 724), TO03 has three times more significantly up-regulated and ten times more significantly down-regulated proteins (Fig. 1, overview Fig. S 5 and S 6). For this reason, if not noted otherwise, we will focus on the TO03 results only (Table 2, Table 3). A short discussion on the different adaptational strategies of the two strains can be found in Notes S 1. The data for all relevant proteins in both strains, and both proteomic datasets (main and extended) are given in the Supplementary Table S 2 – S 9. Expression differences are classed as “highly regulated” (greater than or equal to 2-fold difference) or “regulated” (1.3 to 2-fold difference, see Methods). All differential expression data discussed in the text are significantly up- or down-regulated (*p*<0.05), unless otherwise noted.

**Fig. 1:**
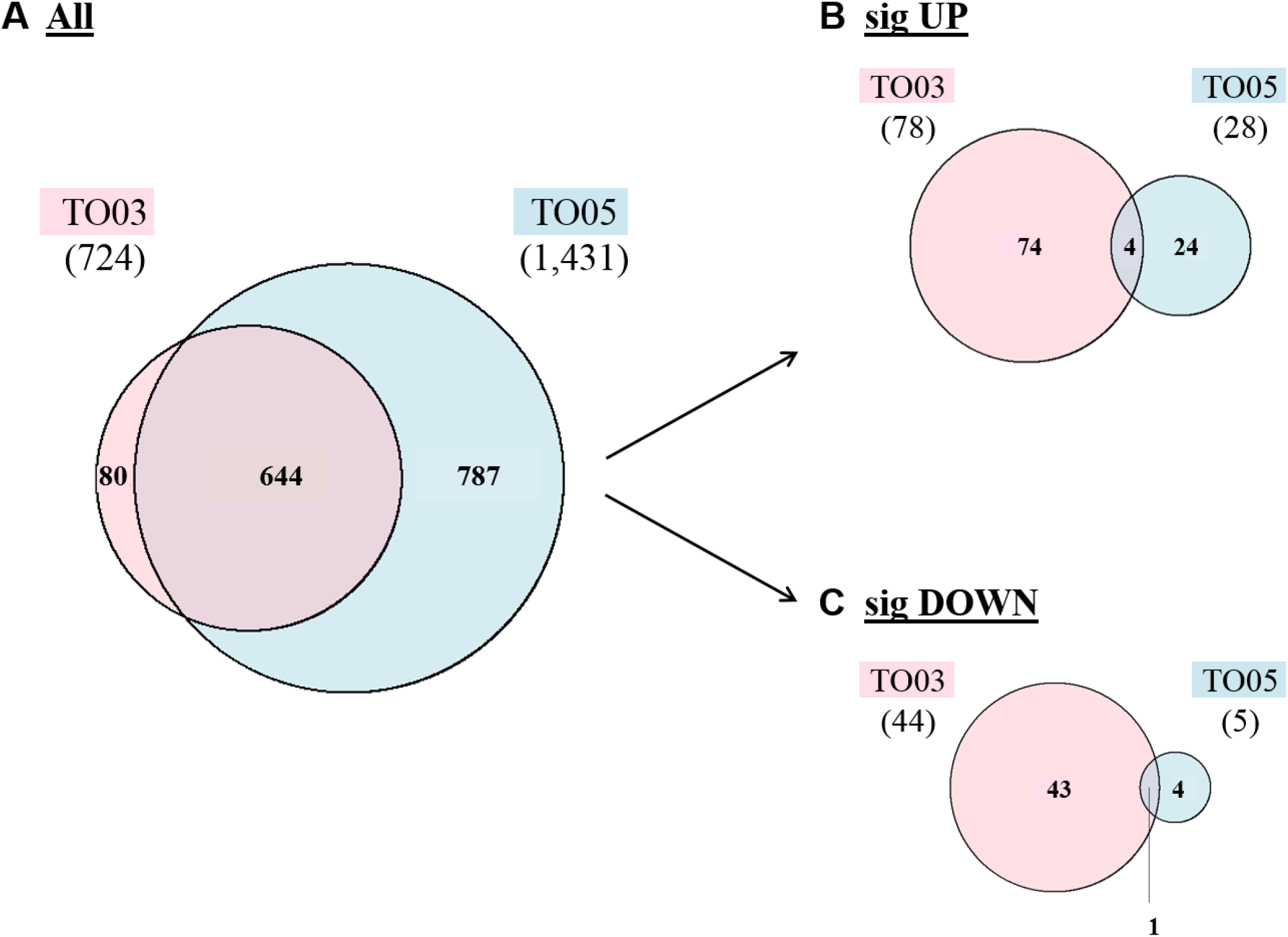
Overview of proteomics data for *T. oceanica* CCMP 1003 (TO03) and 1005 (TO05) grown in Cu-limiting conditions. A-C Venn diagrams of distinct identified proteins in the original datasets of TO03 and TO05. A) All proteins; B) significantly up-regulated proteins; C) significantly down-regulated proteins. Note that even though only 50% of the proteins identified in TO05 were identified in TO03, in TO03 there were three times more significantly up-regulated proteins and five times more significantly down-regulated proteins than in TO05.

**Table 1:**
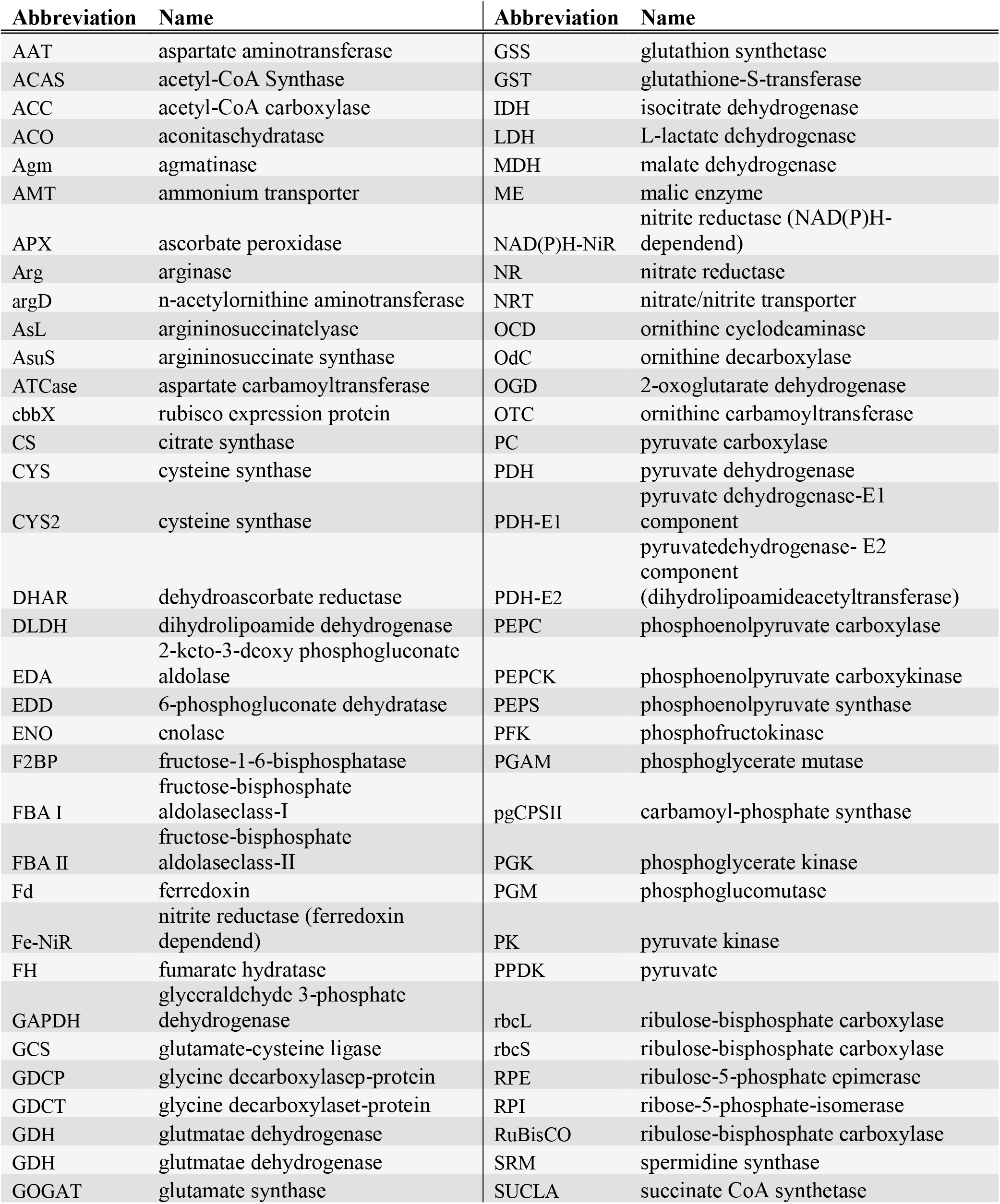
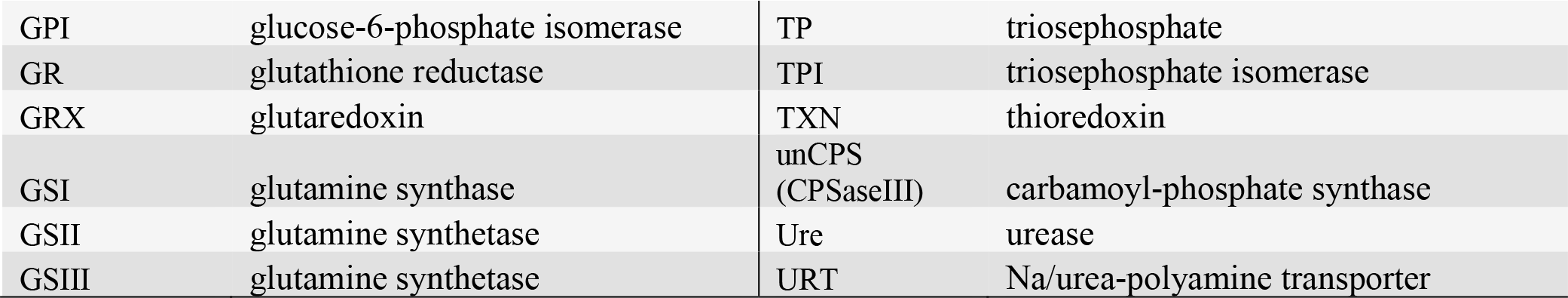
Abbreviations of proteins discussed in this paper.

**Table 2:**
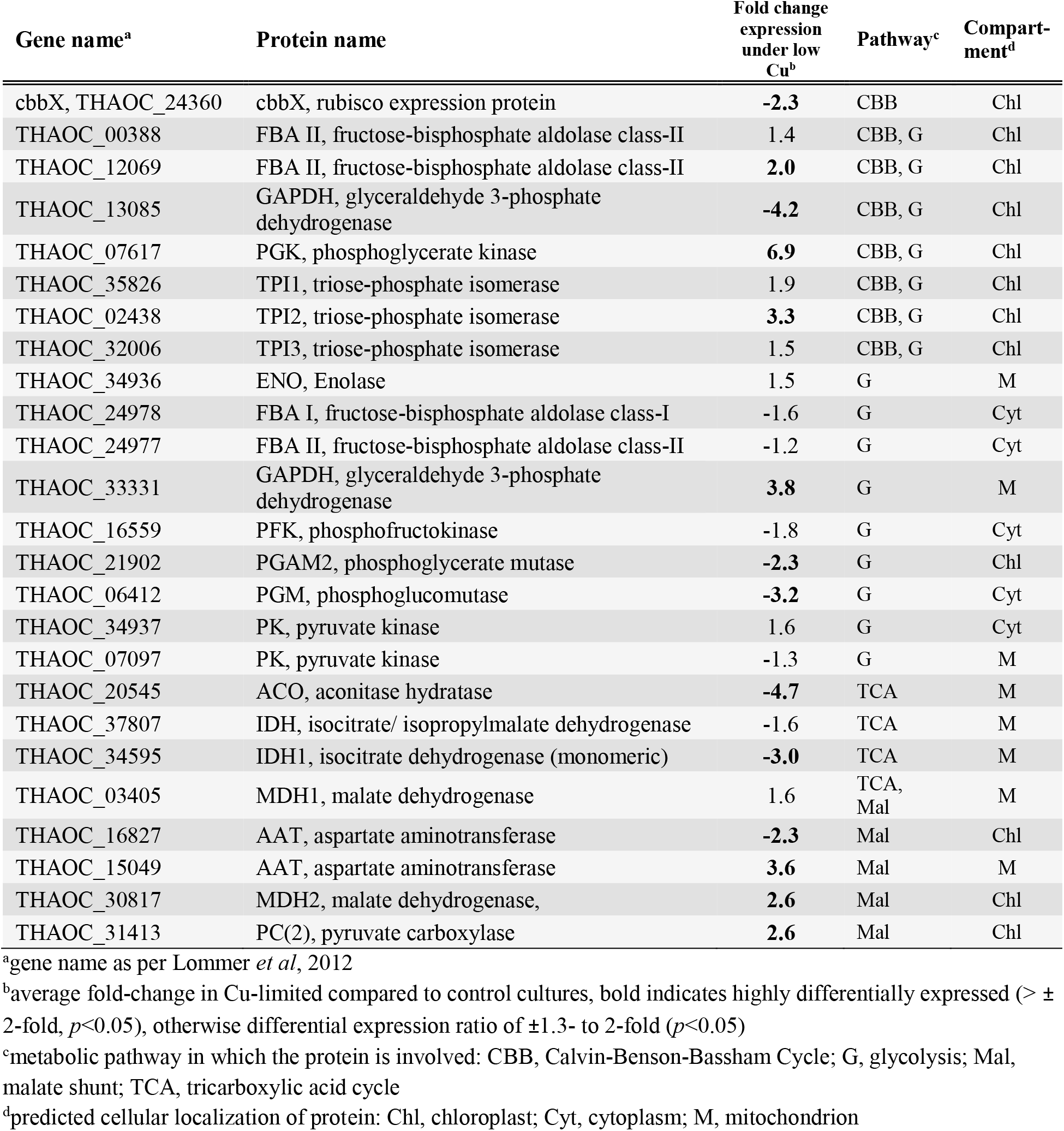
Proteins involved in carbon metabolism that are significantly up- or down-regulated in *T. oceanica* (CCMP 1003) under chronic Cu limitation.

| Gene name<sup>a</sup> | Protein name | Fold change expression under low Cu<sup>b</sup> | Pathway<sup>c</sup> | Compartment<sup>d</sup> |
| --- | --- | --- | --- | --- |
| cbbX, THAOC_24360 | cbbX, rubisco expression protein | **-2.3** | CBB | Chl |
| THAOC_00388 | FBA II, fructose-bisphosphate aldolase class-II | 1.4 | CBB, G | Chl |
| THAOC_12069 | FBA II, fructose-bisphosphate aldolase class-II | **2.0** | CBB, G | Chl |
| THAOC_13085 | GAPDH, glyceraldehyde 3-phosphate dehydrogenase | **-4.2** | CBB, G | Chl |
| THAOC_07617 | PGK, phosphoglycerate kinase | **6.9** | CBB, G | Chl |
| THAOC_35826 | TPI1, triose-phosphate isomerase | 1.9 | CBB, G | Chl |
| THAOC_02438 | TPI2, triose-phosphate isomerase | **3.3** | CBB, G | Chl |
| THAOC_32006 | TPI3, triose-phosphate isomerase | 1.5 | CBB, G | Chl |
| THAOC_34936 | ENO, Enolase | 1.5 | G | M |
| THAOC_24978 | FBA I, fructose-bisphosphate aldolase class-I | -1.6 | G | Cyt |
| THAOC_24977 | FBA II, fructose-bisphosphate aldolase class-II | -1.2 | G | Cyt |
| THAOC_33331 | GAPDH, glyceraldehyde 3-phosphate dehydrogenase | **3.8** | G | M |
| THAOC_16559 | PFK, phosphofructokinase | -1.8 | G | Cyt |
| THAOC_21902 | PGAM2, phosphoglycerate mutase | **-2.3** | G | Chl |
| THAOC_06412 | PGM, phosphoglucomutase | **-3.2** | G | Cyt |
| THAOC_34937 | PK, pyruvate kinase | 1.6 | G | Cyt |
| THAOC_07097 | PK, pyruvate kinase | -1.3 | G | M |
| THAOC_20545 | ACO, aconitase hydratase | **-4.7** | TCA | M |
| THAOC_37807 | IDH, isocitrate/ isopropylmalate dehydrogenase | -1.6 | TCA | M |
| THAOC_34595 | IDH1, isocitrate dehydrogenase (monomeric) | **-3.0** | TCA | M |
| THAOC_03405 | MDH1, malate dehydrogenase | 1.6 | TCA, Mal | M |
| THAOC_16827 | AAT, aspartate aminotransferase | **-2.3** | Mal | Chl |
| THAOC_15049 | AAT, aspartate aminotransferase | **3.6** | Mal | M |
| THAOC_30817 | MDH2, malate dehydrogenase, | **2.6** | Mal | Chl |
| THAOC_31413 | PC(2), pyruvate carboxylase | **2.6** | Mal | Chl |

<sup>a</sup>gene name as per Lommer *et al*, 2012
<sup>b</sup>average fold-change in Cu-limited compared to control cultures, bold indicates highly differentially expressed ( $> \pm 2$ -fold, $p < 0.05$ ), otherwise differential expression ratio of $\pm 1.3$ - to 2-fold ( $p < 0.05$ )
<sup>c</sup>metabolic pathway in which the protein is involved: CBB, Calvin-Benson-Bassham Cycle; G, glycolysis; Mal, malate shunt; TCA, tricarboxylic acid cycle
<sup>d</sup>predicted cellular localization of protein: Chl, chloroplast; Cyt, cytoplasm; M, mitochondrion

**Table 3:** Proteins involved in nitrogen and stress response metabolism that are significantly up- or down-regulated in *T. oceanica* (CCMP 1003) under chronic Cu limitation.

| Gene name<sup>a</sup> | Protein name | fold change expression under low Cu<sup>b</sup> | Pathway<sup>c</sup> | Compartment<sup>d</sup> |
| --- | --- | --- | --- | --- |
| THAOC_15049 | AAT, aspartate aminotransferase | **3.6** | N | M |
| THAOC_16827 | AAT, aspartate aminotransferase | **-2.3** | N | Chl |
| THAOC_02363 | Fe-NiR, nitrite/sulfite reductase ferredoxin-like half-domain | **2.3** | N | Chl |
| THAOC_00016 | Fe-NiR, nitrite/sulfite reductase ferredoxin-like half-domain | **1.3** | N | Chl |
| THAOC_31900 | GSII, glutamine synthetase | 1.7 | N | Chl |
| THAOC_06032 | GSIII, glutamine synthetase | **-5.3** | N | M |
| THAOC_34460 | NR, nitrate reductase | 1.5 | N | Cyt |
| THAOC_00016 | NR, nitrite reductase | 1.3 | N | Chl |
| THAOC_04919 | NRT, nitrate/nitrite transporter | **11.0** | N | Trans |
| THAOC_04380 | OCD, ornithine cyclodeaminase | 1.2 | N | Cyt |
| THAOC_05385 | OTC, ornithine carbamoyltransferase | 1.7 | N | M |
| THAOC_22108 | SRM, spermidine synthase | **2.3** | N | Cyt |
| THAOC_31656 | URT, Na/urea-polyamine transporter | **6.9** | N | Trans |
| THAOC_13288 | GOGAT, glutamate synthase | 1.6 | N, ROS | Chl |
| THAOC_37364 | APX, ascorbate peroxidase | 1.4 | ROS | Cyt |
| THAOC_27524 | CYS, cysteine synthase | **2.5** | ROS | Chl |
| THAOC_10442 | CYS2, cysteine synthase | 1.5 | ROS | Chl |
| THAOC_07268 | GR, glutathione reductase | **2.5** | ROS | Chl |
| THAOC_07269 | GRX, glutaredoxin | -1.3 | ROS | Chl |
| THAOC_18234 | GRX, glutaredoxin | -1.3 | ROS | Chl |
| THAOC_09062 | GST, glutathione-S-transferase | **7.3** | ROS | Cyt |
| THAOC_02860 | MnSOD, Mn/Fe binding superoxide dismutase | 1.8 | ROS | Chl |
| THAOC_10112 | NiSOD, nickel-dependent superoxide dismutase | 1.4 | ROS | Cyt |
| THAOC_05213 | TXN, thioredoxin | **4.2** | ROS | Cyt |
| THAOC_13865 | TXN, thioredoxin | 1.8 | ROS | M |
| THAOC_31425 | TXN, thioredoxin | 1.5 | ROS | Chl |
| THAOC_25559 | Fdx, ferredoxin | **43.79** | --- | Chl |

<sup>a</sup>gene name as per Lommer *et al*, 2012
<sup>b</sup>average fold-change in Cu-limited compared to control cultures, bold indicates highly differentially expressed ( $> \pm 2$ -fold, $p < 0.05$ ), otherwise differential expression ratio of $\pm 1.3$ - to 2-fold ( $p < 0.05$ )
<sup>c</sup>metabolic pathway in which the protein is involved: N, nitrogen metabolism; ROS, reactive oxygen species metabolism; M, multiple
<sup>d</sup>predicted cellular localization of protein: Chl, chloroplast; Cyt, cytoplasm; M, mitochondrion; TRANS, transmembrane

Of the 724 distinctive proteins in TO03, 525 have associated Kegg Orthology (KO) identifiers, and 52% of these were related to metabolism (Fig. 2). Furthermore, 77-78% of these metabolic proteins were particularly affected by Cu limitation, with general trends of down-regulation of proteins involved in energy metabolism, up-regulation of those in carbohydrate metabolism, and a modification of those in amino acid metabolism.

**Fig. 2:**
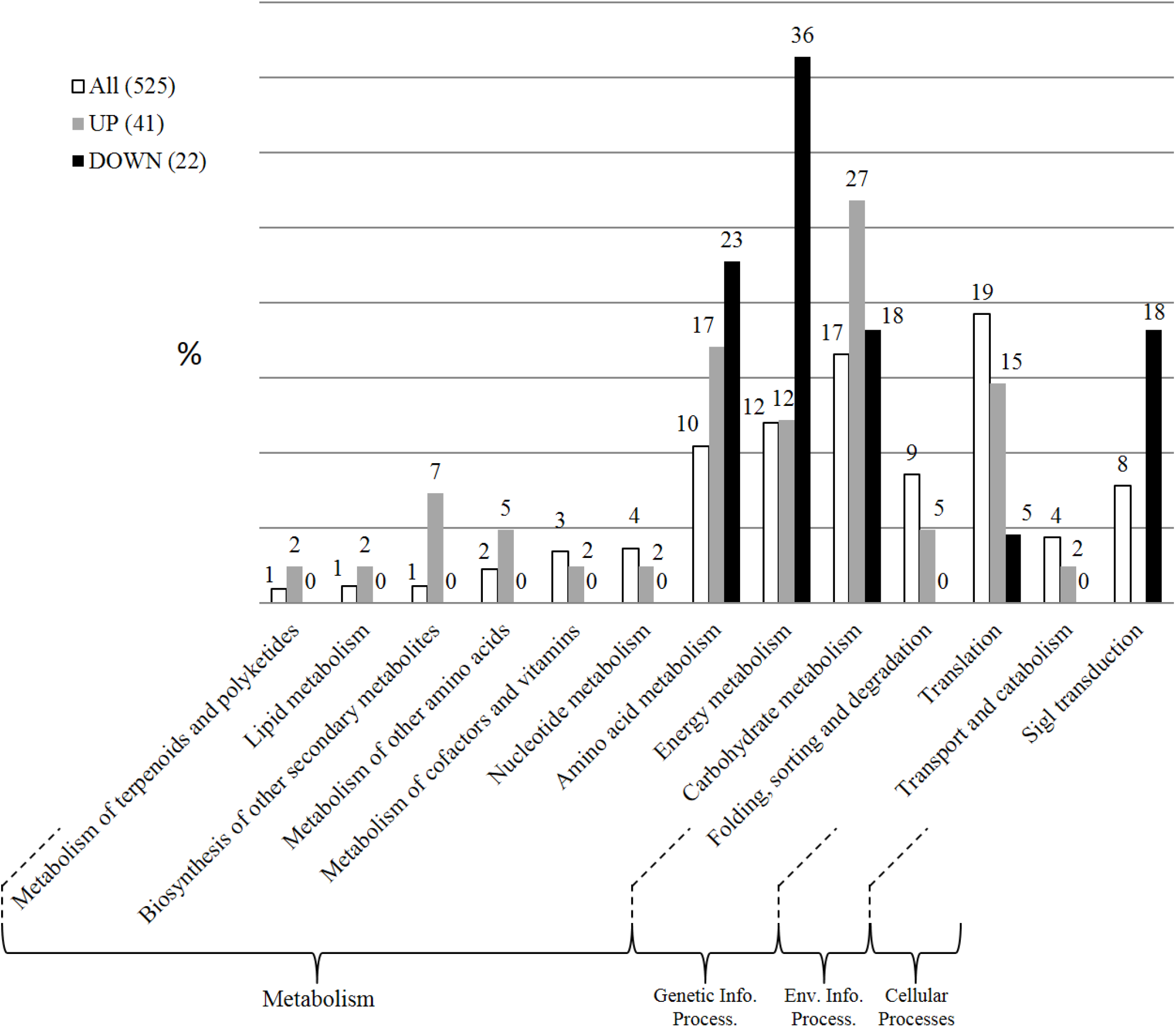
Bar charts of second level Kegg Orthology (KO) identifiers associated with proteins of original TO03 dataset comparing proportions between all proteins and those that are significantly up- or down-regulated. Numbers in brackets indicate absolute protein numbers in each set. Note that certain aspects of metabolism are most highly affected: energy, amino acid and carbohydrate metabolism

### Carbon fixation, Glycolysis and the Citrate (TCA) cycle

In diatoms the enzymes of glycolysis are found in all three major compartments: chloroplast stroma, cytosol and mitochondria (Gruber and Kroth, 2017; Kroth et al., 2008; Río Bártulos et al., 2018; Smith et al., 2012). Four (or seven, if the 3 triose isomerase isoenzymes are counted) of the 15 proteins involved in the carbon fixing Calvin-Benson-Bassham (CBB) cycle are part of the chloroplast glycolytic pathway (Table 2, Fig. 3 (CBB and TCA cycle), Fig. 4 (Glycolysis), Table S 4). In the initial step of CO2 fixation, the large and small subunits of Rubisco were not affected by Cu limitation but the essential Rubisco activator protein cbbX (To24360) was down-regulated by 2.3-fold. Six proteins were up-regulated: phosphoglycerate kinase (PGK, To07617) by 6.8-fold, the two fructose-bisphosphate aldolase, class II proteins by 1.4 and 2-fold (FBA II, To00388 and To12069), and the three triose phosphate isomerase isoenzymes (TPI, To02438, To35826, To32006) by 3.3-, 1.9- and 1.5-fold, respectively. Glyceraldehyde 3-phosphate dehydrogenase (GAPDH, To13085) and phosphoglycerate mutase (PGAM, To21902, 2.3-fold) were the only proteins down-regulated, by 4.2- and 2.3-fold, respectively. Of the nine expressed proteins targeted to the chloroplast, only fructose-bisphosphate aldolase, class I (FBA I, To02112) was not affected by Cu limitation.

**Fig. 3:**
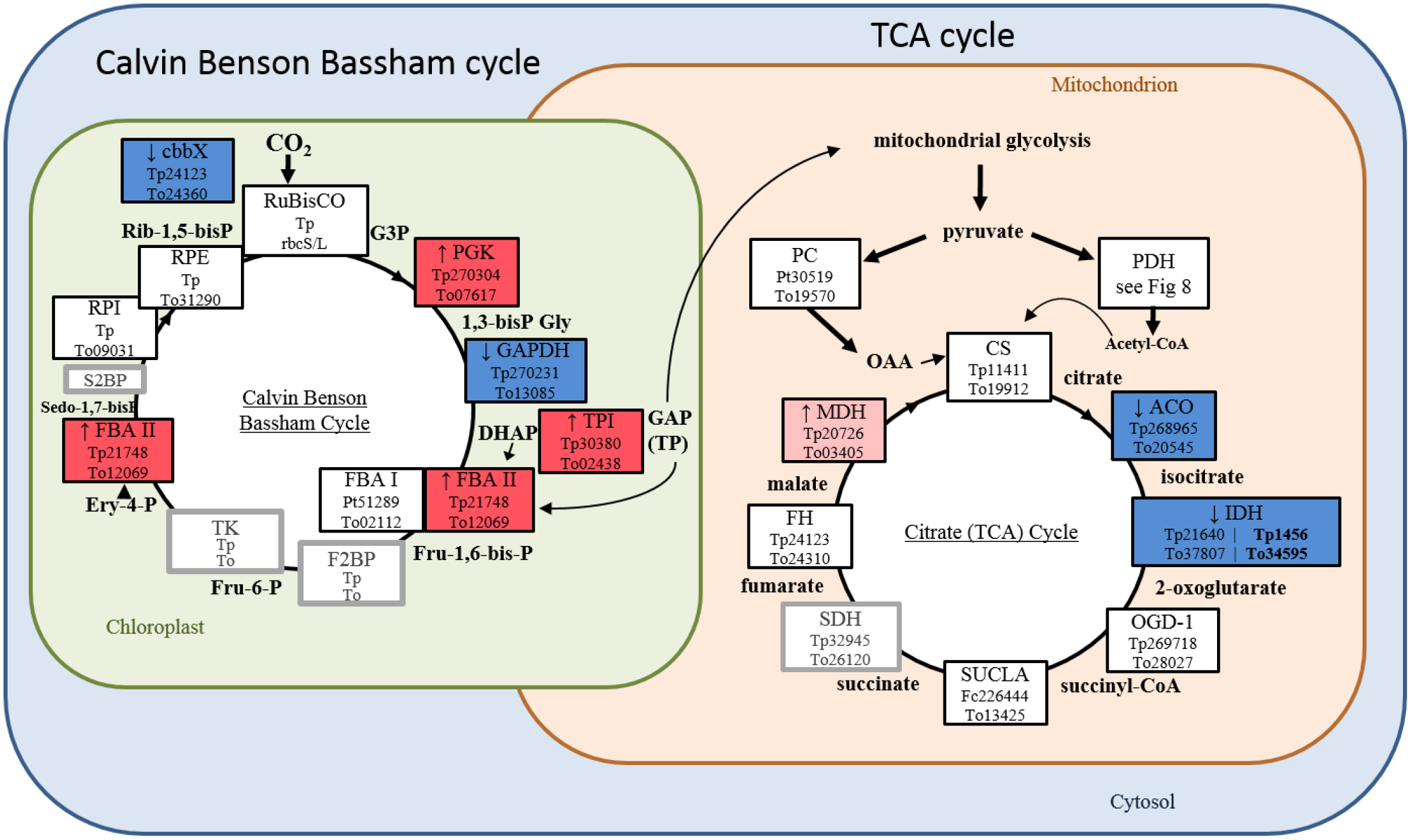
Relative expression of proteins involved in the Calvin Benson Bassham and citrate (TCA) cycle. Boxes indicate proteins with their abbreviated name and known *T. pseudonana* (Tp) and *T. oceanica* (To) homologs. The colors of the boxes indicate expression in *Toceanica* TO03 under low Cu: dark red, highly up-regulated (>2-fold, *p*<0.05); light pink, up-regulated by 1.3 to 2-fold (*p*<0.05); dark blue, highly down-regulated (>2-fold, *p*<0.05); light blue, down-regulated by 1.3 to 2-fold (*p*<0.05); white, expressed in TO03; grey border around box, found in *T.oceanica* T005 genome but not expressed in TO03 proteomic data; grey, dashed border around box, no putative homologs in the *T. oceanica* genome. **Protein abbreviations:** ACO, aconitase hydratase; cbbX, rubisco expression protein; CS, citrate synthase; DLDH, dihydrolipoamide dehydrogenase; FBA I, fructose-bisphosphate aldolase class-I; FBA II, fructose-bisphosphate aldolase class-II; FH, fumarate hydratase; GAPDH, glyceraldehyde 3-phosphate dehydrogenase; GPI, glucose-6-phosphate isomerase; IDH, isocitrate dehydrogenase; MDH, malate dehydrogenase; OGD, 2-oxoglutarate dehydrogenase; PC, pyruvate carboxylase; PDH, pyruvate dehydrogenase; PFK, phosphofructokinase; PGAM, phosphoglycerate mutase; PGK, phosphoglycerate kinase; PGM, phosphoglucomutasePK, pyruvate kinase; rbcL, ribulose-bisphosphate carboxylase large chain; rbcS, ribulose-bisphosphate carboxylase small chain; RPE, ribulose-5-phosphate epimerase; RPI, ribose-5-Phosphate-isomerase; SUCLA, succinate CoA synthetase; TPI, triose-phosphate isomerase **Compound abbreviations**: 1,3-bisPG, 1,3-bisphosphateglycerate; G3P, glucose-3-phosphate; Ery-4-P, erythrose 4 phosphate; Sedo-1,7-bisP, sedoheptulose 1 7-bisphosphate; Rib-1,5-bisP, ribulose-1,5-bisphosphate; DHAP, dihydroxyacetone phosphate; Fru-1,6-bis-P, fructose 1,6-bisphosphate; Fru-6-P, fructose 6-phosphate; GAP, glyceraldehyde 3-phosphate; HCO_3_^-^, bicarbonate; OAA, oxaloacetate; TP, triose phosphate

**Fig. 4:**
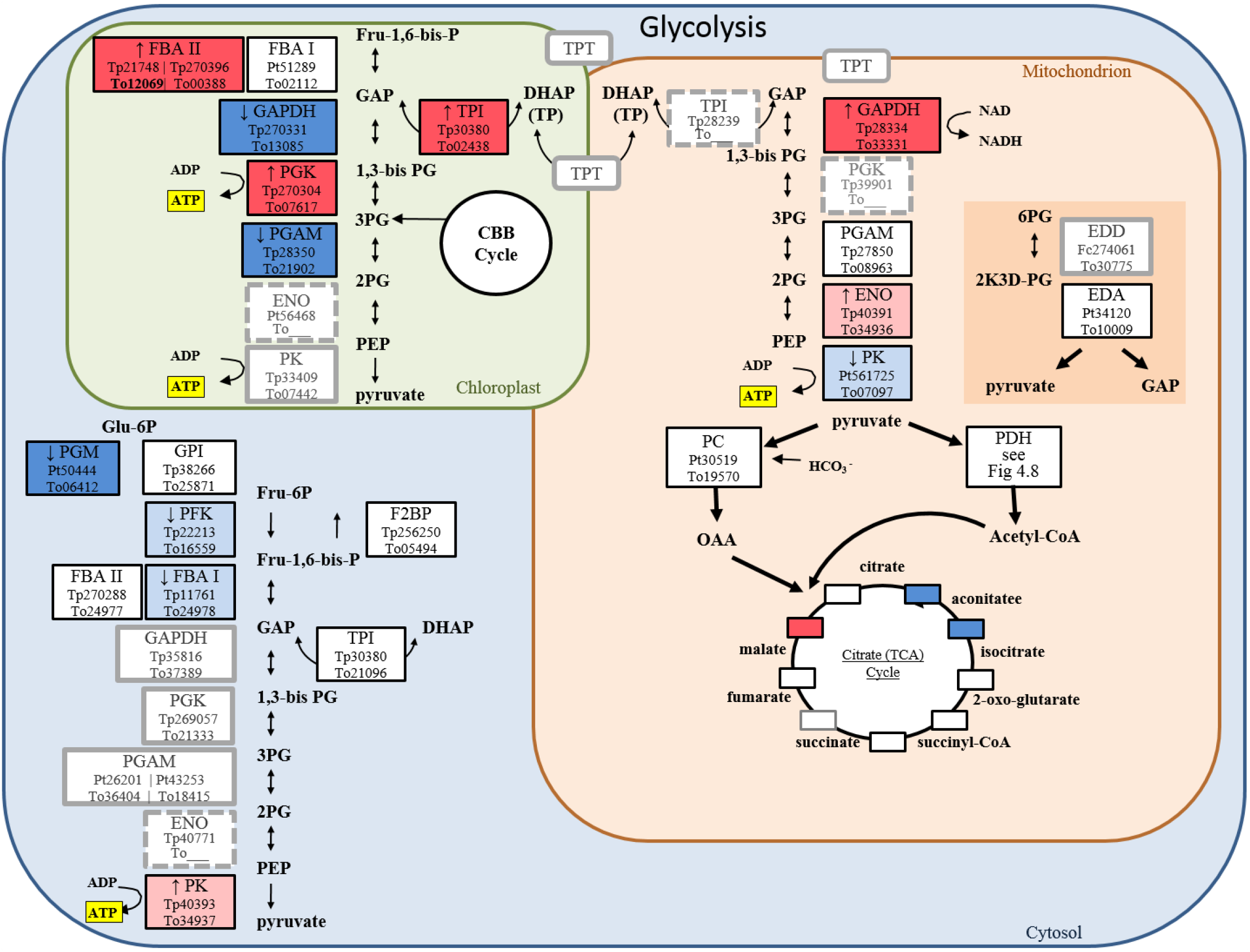
Relative expression of proteins involved in glycolysis in the three compartments (chloroplast, mitochondrion, cytosol), including Entner-Dourdoroff pathway. Boxes indicate proteins with their abbreviated name and known *T. pseudonana* (Tp) and *T. oceanica* (To) homologs. The colors of the boxes indicate expression in *Toceanica* TO03 under low Cu: dark red, highly up-regulated (>2-fold, *p*<0.05); light pink, up-regulated by 1.3 to 2-fold (*p*<0.05); dark blue, highly down-regulated (>2-fold, *p*<0.05); light blue, down-regulated by 1.3 to 2-fold (*p*<0.05); white, expressed in TO03; grey border around box, found in *T. oceanica* T005 genome but not expressed in TO03 proteomic data; grey, dashed border around box, no putative homologs in the *T. oceanica* genome. **Protein abbreviations**: CBB cycle, Calvin Benson Bassham cycle, EDA, 2-keto-3-deoxy phosphogluconate aldolase; EDD, 6-phosphogluconate dehydratase; ENO, enolase; F2BP, fructose-1-6-bisphosphatase; FBA I, fructose-bisphosphate aldolase class-I; FBA II, fructose-bisphosphate aldolase class-II; GAPDH, glyceraldehyde 3-phosphate dehydrogenase; GPI, glucose-6-phosphate isomerase; PC, pyruvate carboxylase; PDH, pyruvate dehydrogenase; PFK, phosphofructokinase; PGAM, phosphoglycerate mutase; PGK, phosphoglycerate kinase; PGM, phosphoglucomutase; PK, pyruvate kinase; TCA, tricarboxylic acid cycle;TP, triose phosphate; TPI, tiose-phosphate isomerase, TPT, triose phosphate transporter **Compound abbreviations**: 1,3-bisPG, 1,3-bisphosphateglycerate; 2K3D-PG, 2-keto-3-deoxyphosphogluconate; 2PG, 2-phosphoglycerate; 3PG, 3-phosphoglycerate; 6PG, 6-phosphogluconate; DHAP, dihydroxyacetone phosphate; Fru-1,6-bis-P, fructose 1,6-bisphosphate; Fru-6P, fructose 6-phosphate; GAP, glyceraldehyde 3-phosphate; Glu 6-P, glucose 6-phosphate; HCO_3_^-^, bicarbonate; OAA, oxaloacetate; PEP, phosphoenolpyruvate

The up-regulation of TPI (Fig. 3, Fig. 4) combined with the down-regulation of GAPDH and PGAM could lead to an increase in triose-phosphates and their subsequent export from the chloroplast. Probing the genome for gene models containing the triose-phosphate transporter Pfam domain identified seven candidate genes (Table S 3) of which only two were expressed. Neither of them was differentially expressed.

Nine expressed proteins involved in the citrate cycle in the mitochondria were identified (Fig. 3, Table 2, Table S 5). Malate dehydrogenase (MDH1, To03405) was the only one up-regulated (1.6-fold). Aconitase hydratase (ACO, To20545) and two isocitrate dehydrogenases (IDH, To37807, To34595) were all down-regulated by 4.7-, 3.0-, and 1.6-fold, respectively. Of the proteins considered to be part of mitochondrial glycolysis (Fig. 4), glyceraldehyde-3-phosphate dehydrogenase (GAPDH, To33331) and enolase (ENO, To34936) were both up-regulated by 3.8 and 1.5-fold respectively, while pyruvate kinase (PK, To07097) was down-regulated by 1.3-fold.

Of the eight expressed cytosolic proteins detected, three were down-regulated: phosphoglucomutase (PGM, To06412) by 3.2-fold, phosphofructokinase (PFK, To16559) by 1.8-fold, and fructose-bisphosphate aldolase, class I (FBA I, To24978) by 1.6-fold (or 2.7-fold considering expression of a contig associated with the same gene). The only cytosolic protein that was up-regulated was pyruvate kinase (PK, To34937, by 1.6-fold).

### Nitrogen metabolism

Twenty-two proteins involved in the urea cycle, nitrogen acquisition and assimilation, as well as four membrane transporters were identified (Table 3, Fig. 5, Table S 6). At the plasma membrane, the urea (URT, To31656) and nitrate/nitrite (NRT, To04919) transporters were both significantly up-regulated (6.9 and 11-fold, respectively). However, the expression of the two transporters putatively located in the chloroplast envelope, the formate/nitrate (NiRT, To00240) and ammonium (AMT, To07247) transporters were not affected by Cu limitation.

**Fig. 5:**
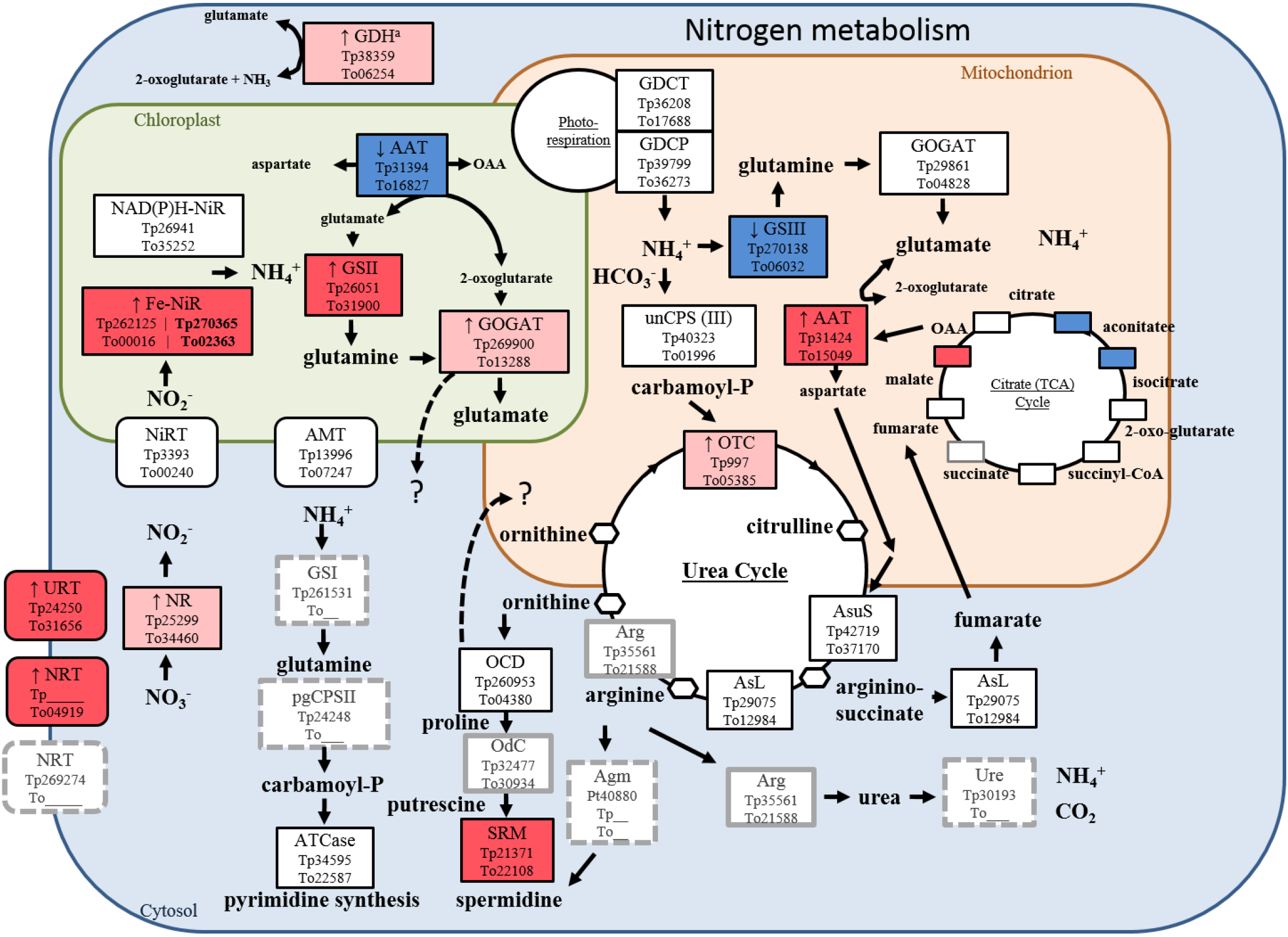
Relative expression of proteins involved in nitrogen metabolism. Boxes indicate proteins with their abbreviated name and known *T. pseudonana* (Tp) and *T. oceanica* (To) homologs. The colors of the boxes indicate expression in *Toceanica* TO03 under low Cu: dark red, highly up-regulated (>2-fold, *p*<0.05); light pink, up-regulated by 1.3 to 2-fold (*p*<0.05); dark blue, highly down-regulated (>2-fold, *p*<0.05); light blue, down-regulated by 1.3 to 2-fold (*p*<0.05); white, expressed in TO03; grey border around box, found in *T. oceanica* T005 genome but not expressed in TO03 proteomic data; grey, dashed border around box, no putative homologs in the *T. oceanica* genome. **Protein abbreviations**: AAT, aspartate aminotransferase; Agm, agmatinase; AMT, ammonium transporter; Arg, arginase; argD, n-acetylornithine aminotransferase; AsL, argininosuccinate lyase; AsuS, argininosuccinate synthase; ATCase, aspartate carbamoyltransferase; Fe-NiR, nitrite reductase (ferredoxin-dependent); GDCP, glycine decarboxylase p-protein; GDCT, glycine decarboxylase t-protein; GDH, glutamate dehydrogenase; GOGAT, glutamate synthase; GSI, glutamine synthase; GSII, glutamine synthetase; GSIII, glutamine synthetase; NAD(P)H-NiR, nitrite reductase (NAD(P)H dependent); NiRT, formate/nitrite transporter; NR, nitrate reductase; NRT, nitrate/nitrite transporter; OCD, ornithine cyclodeaminase; OdC, ornithine decarboxylase; OTC, ornithine carbamoyltransferase; pgCPSII, carbamoyl-phosphate synthase II; SRM, spermidine synthase; unCPS (CPSase III), carbamoyl-phosphate synthase; Ure, urease; URT, Na/urea-polyamine transporter.

Within the chloroplast, three nitrite reductases were identified. Of these, the NAD(P)H-dependent isoenzyme was not differentially expressed (NAD(P)H-NiR, To35252), whereas two ferredoxin-containing nitrite reductases (Fe-NiR, To00016 and To02363) were up-regulated by 1.3- and 2.3-fold, respectively, in concert with the massive 40-fold increase of ferredoxin in the chloroplast (Table 3; Hippmann et al., 2017). Glutamine (GSII, To31900) and glutamate synthases (GOGAT, To13288) which both require ferredoxin cofactors were both up-regulated (1.7- and 1.6-fold, respectively). In contrast, aspartate aminotransferase (AAT, To16827) was the only chloroplast protein involved in core nitrogen metabolism that was down-regulated (2.3-fold).

In the mitochondria, the pattern was reversed: the glutamine synthase isoform (GSIII, To06032) was 5.3-fold down-regulated, while the glutamate synthase isoform (GOGAT, TO04828) expression did not change, and the mitochondrial aspartate aminotransferase (AAT, To15049) was up-regulated by 3.6-fold. Glycine decarboxylase t- and p-proteins (GDCT/P, To17688 and To36273), involved in photorespiration, were not affected. In the urea cycle, six proteins were identified but only ornithine carbamoyltransferase (OTC, To05385) was up-regulated by 1.7-fold.

In the cytosol, glutamate dehydrogenase (GDH, To06254) was up-regulated by 2.09-fold (p = 0.05) and nitrate reductase (NR, To34460) was up-regulated by 1.5-fold. The only other cytosolic protein upregulated was spermidine synthase (SRM, To22108; by 2.3-fold), which is essential for silica deposition.

### Malate shunt

In plants, malate transfers excess NAD(P)H reducing equivalents from one compartment (i.e. the chloroplast) to another (i.e. the mitochondria) (reviewed by (Scheibe, 2004)). Metabolite antiporters and two isoenzymes for malate dehydrogenase (MDH) and amino aspartate transaminase (AAT) are involved (Table 2, Fig. 6, Table S 8). In diatoms, the malate shunt has been proposed to connect the chloroplast with the mitochondria (Bailleul et al., 2015; Prihoda et al., 2012). In *P. tricornutum*, both MDH1 and MDH2 are targeted to the mitochondrion (Ewe et al., 2018) but in *T. pseudonana* MDH2 is predicted to be targeted to the chloroplast (Smith et al., 2012). Aligning the *T. oceanica* model with Smith’s extended TpMDH2 model (Fig. S 2), supports the conclusion that ToMDH2 is also targeted to the chloroplast.

**Fig. 6:**
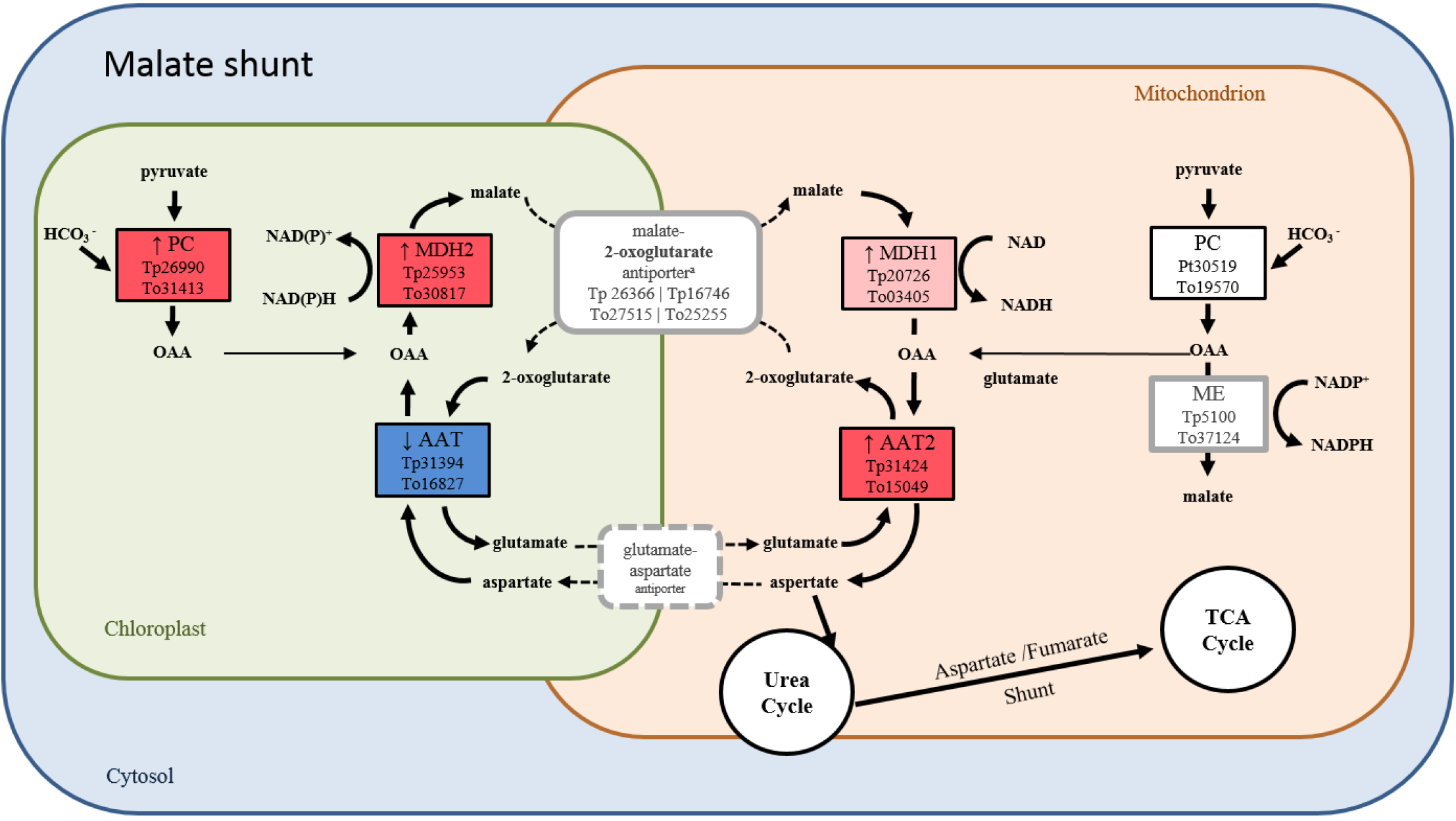
Relative expression of proteins involved in the malate shunt. Boxes indicate proteins with their abbreviated name and known *T. pseudonana* (Tp) and *T. oceanica* (To) homologs. The colors of the boxes indicate expression in *Toceanica* TO03 under low Cu: dark red, highly up-regulated (>2-fold, *p*<0.05); light pink, up-regulated by 1.3 to 2-fold (*p*<0.05); dark blue, highly down-regulated (>2-fold, *p*<0.05); light blue, down-regulated by 1.3 to 2-fold (*p*<0.05); white, expressed in TO03; grey border around box, found in *T. oceanica* T005 genome but not expressed in TO03 proteomic data; grey, dashed border around box, no putative homologs in the *T. oceanica* genome. **Abbreviations**: AAT, aspartate aminotransferase; MDH, malate dehydrogenase; ME, malic enzyme; OAA, oxaloacetate; PC, pyruvate carboxylase; PK, pyruvate kinase

In our proteomic datasets, we found evidence for reciprocal regulation of both isonzyme sets in the putative malate shunt (MDH1, MDH2, AAT, AAT2), as well as two isoenzymes of pyruvate carboxylase (PC) that could be feeding into this metabolic pathway (Fig. 6). Of these 6 enzymes, 4 were up-regulated: both, plastidial pyruvate carboxylase (PC, To31413) and malate dehydrogenase (MDH2, To30817) by 2.6-fold and mitochondrial MDH1 (To03405) and AAT2 (To15049) by 1.6- and 3.6-fold, respectively. Only the plastidial AAT (To16827) was down-regulated (by 2.3-fold).

### Glutathione and antioxidant metabolism strongly upregulated

Glutathione is a small tripeptide (Glu-Cys-Gly) that is involved in redox sensing and counteracting ROS. Twenty-one expressed proteins involved in glutathione metabolism and other antioxidant agents (eg. three thioredoxins, three glutaredoxins, and three superoxide dismutases) were identified (Table 3, Fig. 7, Table S 7). Nine proteins are predicted to be targeted to the chloroplast. Six of these were up-regulated: two isoenzymes for cysteine synthase (CYS, To27524 by 2.5-fold, To10442 by 1.5-fold), glutamate synthase (GOGAT, To13288 by 1.6-fold), glutathione reductase (GR, To07268, by 2.5-fold), thioredoxin (TXN, To31425, by 1.5-fold), and the Mn-containing SOD (MnSOD, To02860, by 1.8-fold). Two glutaredoxin isoenzymes (GRX; To07269, To18234) were only mildly down-regulated proteins (both by 1.3-fold).

**Fig. 7:**
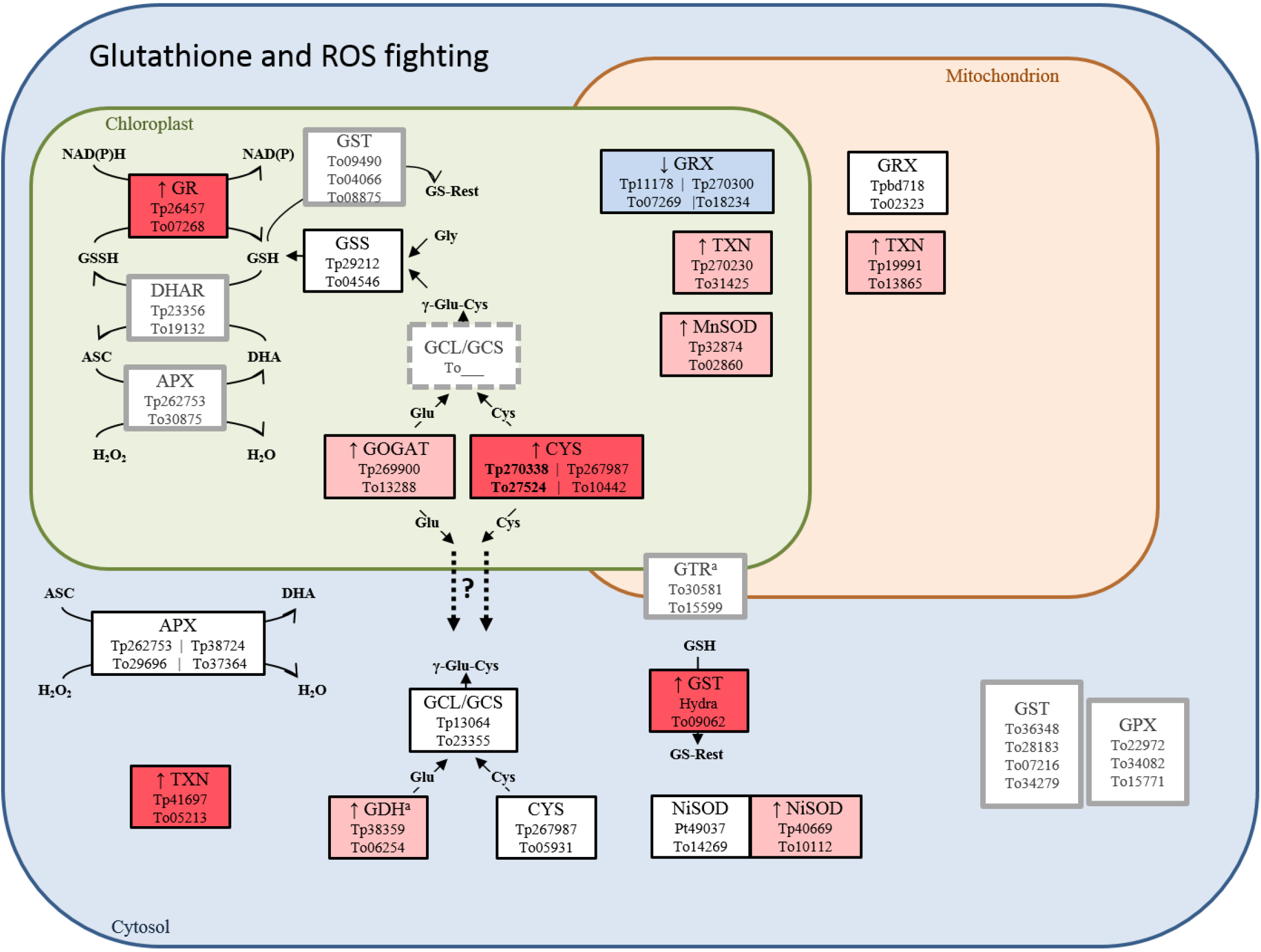
Relative expression of proteins involved in glutathione metabolism and response to reactive oxygen species (ROS). Boxes indicate proteins with their abbreviated name and known *T. pseudonana* (Tp) and *T. oceanica* (To) homologs. The colors of the boxes indicate expression in *Toceanica* TO03 under low Cu: dark red, highly up-regulated (>2-fold, *p*<0.05); light pink, up-regulated by 1.3 to 2-fold (*p*<0.05); dark blue, highly down-regulated (>2-fold, *p*<0.05); light blue, down-regulated by 1.3 to 2-fold (*p*<0.05); white, expressed in TO03; grey border around box, found in *T. oceanica* T005 genome but not expressed in TO03 proteomic data; grey, dashed border around box, no putative homologs in the *T. oceanica* genome. **Protein and compound abbreviations**: APX, ascorbate peroxidase; Cys, cysteine; CYS, cysteine synthase; DHAR, dehydroascorbate reductase; γ-glu-cys, γ-glutamylcysteine; GCL, glutamate cysteine ligase; GDH, glutamate dehydrogenase, glu, glutamate; NADP dependent; GOGAT, glutamate synthase; GR, glutathione reductase; GRX, glutaredoxin; GSS, glutathione synthetase; GTR, glutathione transporter; TXN, thioredoxin

Of the nine cytosolic proteins, glutathione-S-transferase (GST, To09062) and TXN (To05213) were highly up-regulated (by 7.3- and 4.2-fold, respectively), while one of the two Ni-dependent SOD isoenzymes was moderately up-regulated (NiSOD, To10112, by 1.4-fold). Glutamate-cysteine ligase (GCL/GCS, To23355) was only identified in one of the biological triplicates, but with a 4.5-fold increase in expression. In contrast to the chloroplast CYS isoenzymes, the expression of cytosolic CYS (To05931) did not change. The TO03 GST (To09062) has its closest homologs in the polyp *Hydravulgaris*, the anemone *Nematostella*, and the brachiopod *Lingula*, and not in other diatoms (Table S 7).

Only 2 of the expressed proteins involved in glutathione metabolism are predicted to be targeted to the mitochondria; TXN (To13864) was up-regulated by 1.8-fold, whereas glutaredoxin (GRX, To02323) was not differentially expressed in Cu-limited TO03.

## Discussion

In response to low Cu, *T. oceanica* (CCMP1003) restructures the photosynthetic electron transport proteins, resulting in a decrease in carbon assimilation (mg C mg Chl*a*^-1^ h^-1^), and increased susceptibility to overreduction of the photosynthetic electron transport chain (Hippmann et al., 2017). Susceptibility to over-reduction of the photosynthetic electron transport chain at saturating light intensities was suggested by a) the 17% reduction in photochemical quenching (Fq’/Fv’), and b) the light response curves which indicated that the light saturation point decreased well below the actual growth light of 155 umol quanta m^-2^ s^-1^. Growth at saturating light conditions could therefore lead to an increase in reactive oxygen species (ROS). Consequently, there would be an increased need to safely dissipate excess energy, for example through additional electron sinks (Niyogi, 2000). Our findings of a ~40-fold increase in ferredoxin (Fd, petF) and a 2.5-fold increase in ferredoxin: NAD(P)H oxidoreductase (FNR) under Cu limitation (Hippmann et al. 2017) suggested that there is indeed a surplus of reduced ferredoxin (Fd^red^) and NAD(P)H in the chloroplast.

Here, based on our now expanded proteomic analysis, we hypothesize how the interaction between various metabolic pathways (e.g. nitrogen assimilation, glycolysis, citrate and the urea cycle) and the sophisticated coordination between the chloroplast and the mitochondria may facilitate the re-oxidation of Fd^red^ and NAD(P)H in the chloroplast. Protein abundance alone is not always indicative of protein activity. and where known, we have included information on posttranslational activity modulation. We discuss our results with this in mind while suggesting a plausible restructuring of key metabolic pathways in *T. oceanica* in response to copper limitation.

### Carbon metabolism – the Calvin-Benson-Bassham cycle is down-regulated via its activase, and glycolysis is used to redistribute ATP and NAD(P)H within the cell

The three most thoroughly annotated diatom genomes [*T. pseudonana*, Armbrust et al, 2004; *P. tricornutum*, Bowler et al, 2008; *F. cylindricus*, Mock et al, 2017] revealed many isoenzymes, particularly those involved in C metabolism. Indeed, homologous C metabolism isoenzymes exist among and between these diatoms (Gruber and Kroth, 2017; Kroth et al., 2008; Smith et al., 2012), and their differential expression is thought to manage cellular carbon flow. Furthermore, given that within the chloroplast more than 50% of the proteins involved in glycolysis are also part of the Calvin-Benson-Bassham cycle (Smith et al., 2012), to regulate C flow, some isoenzymes might be preferentially involved in glycolysis over carbon fixation. For example, in *P. tricornutum*, the three plastidial fructose-bisphosphate aldolases (FBAs) are differently targeted and regulated under low *vs*. high Fe conditions (Allen et al., 2012)). Here, we hypothesize that to overcome Cu limitation, *T. oceanica* down-regulates the Calvin-Benson-Bassham cycle, while modulating glycolysis to promote the redistribution of ATP and NAD(P)H reducing equivalents among cellular compartments.

Similarly to *P. tricornutum* under Fe limitation, Cu-limited *T. oceanica* also regulates the expression of FBA homologs (Table 4), albeit in a different way. While the chloroplast FBA (FbaC2 homolog, To12069) is up-regulated, one of the pyrenoid-associated FBAs is only mildly up-regulated (FbaC1 homolog, To00388). This suggests that FbaC2 is preferentially involved in glycolysis over C assimilation, for the following reasons: (1) C assimilation decreased by 66% in Cu-limited cultures compared to the control (Hippmann et al, 2017), suggesting it is less likely for the C fixation proteins to be up-regulated; (2) the three significantly up-regulated proteins involved in the Calvin-Benson-Bassham cycle can also be part of glycolysis (i.e. PGK, TPI, and FBA, Table 2); (3) none of the distinct Calvin-Benson-Bassham cycle proteins (i.e. Rubisco, RPI, and RPE) were differentially expressed; (4) the red algal-type Rubisco activase (cbbX) was down-regulated by 2.25-fold. The down-regulation of cbbX results in slower carbon fixation and activity of Rubisco although Rubisco levels remain unchanged (Mueller-Cajar et al., 2011). Since RPI and RPE abundance remain constant, ribulose-bisphosphate would be bound to Rubisco. Consequently, once nutrient conditions are favorable, only the cbbX would need up-regulation for C fixation to proceed. This strategy might be advantageous in nutrient limited environments with short-lived nutrient-rich conditions.

**Table 4:** Fructose-bisphosphate aldolase (FBA) isoenzymes in *P. tricornutum* (Pt) and homologs in *T. oceanica* (To, CCMP 1003). Information on *P. tricornutum* as per Allen *et al* (2012).

| Gene name (Pt)<sup>a</sup> | FBA class<sup>b</sup> | Phylogenetic ancestry<sup>c</sup> | Location in Pt<sup>d</sup> | Pt id<sup>e</sup> | Pt mRNA lowFe<sup>f</sup> | To homolog<sup>g</sup> | protein ratio lowCu<sup>h</sup> |
| --- | --- | --- | --- | --- | --- | --- | --- |
| FbaC1 | Class II | Chromalveolate specific gene duplication of FbaC2 prior to diversification | Chloroplast, Pyrenoid | Bd825 | ↑ >25 | To00388 | ↑ (1.4 ) |
| FbaC2 | Class II | Endosymbiotic gene transfer from prasinophyte-like green algal ancestor | Chloroplast, diffuse | Pt22993 | ↓ <20 | To12069 | ↑ (2.0 ) |
| Fba3 | Class II | Heterokont host of secondary endosymbiosis | Cytosol | Pt29014 | ↑ >10 | To24977 | ± |
| Fba4 | Class I | Bacterial like (unknown in non-diatom eukaryotes) | Cytosol, putative cytoskeletal interaction | Pt42447 | ~1 | To24978 | ↓ (-2.8 ) |
| FbaC5 | Class I | Endosymbiotic gene transfer from red algal ancestor (with selective gene loss in some centric diatoms) | Chloroplast, Pyrenoid | Pt51289 | ↑ >80 | To02112 | ± |

<sup>a</sup>as per Allen *et al* (2012)
<sup>b</sup>Class I uses a metal co-factor, Class II uses a Schiff base
<sup>c</sup>as per Allen *et al* (2012)
<sup>d</sup>as per Allen *et al* (2012) using GFP-fusion proteins
<sup>e</sup>NCBI identifier
<sup>f</sup>fold-change of mRNA transcript levels in acute Fe limited *vs.* Fe replete cultures; arrows indicating up- and down-regulation
<sup>g</sup>as per blastP search
<sup>h</sup>fold change of protein levels in chronic Cu limited *vs.* Cu replete cultures
FBA, fructose-bisphosphate aldolase; Pt, *Phaeodactylum tricornutum*; To, *Thalassiosira oceanica*

In general, most reactions facilitated by proteins in glycolysis can proceed in either direction, i.e. glycolysis or gluconeogenesis. Smith *et al*. (2012) suggest that gluconeogenesis prevails in the mitochondria. However, assuming that the required metabolite transporters are present in the mitochondria (e.g. aspartate/glutamate shuttle, malate/2–oxoglutarate shuttle, citrate/malate shuttle, and fumarate/succinate shuttle), modelling flux balances in *P. tricornutum* predict that glycolysis would indeed be more favorable than gluconeogenesis in the mitochondria (Kim et al., 2016).

In *T. oceanica*, in each cellular compartment, different subsets of glycolytic proteins were up- or down-regulated under Cu limitation (Fig. 4, Table S 4, overview Fig. S 5). Focusing on the up-regulated proteins (Fig. S 3), a pattern emerges relating the possibility of increased ATP formation in the chloroplast and cytosol with NAD(P)H consumption in the chloroplast and its coupled formation in the mitochondria. By reducing chloroplast GAPDH (To13085) and increasing mitochondrial GAPDH (To33331), NAD(P)H reducing equivalents would be generated in the mitochondria, whereas increasing PGK (To07617) in the chloroplast would increase ATP in this compartment. Therefore, an increased ATP/NAD(P)H ratio in the plastid would be predicted under Cu limitation.

The contrasting differential expression of GAPDH in plastid and mitochondrial compartments suggest the possibility of a key role for triose-phosphate transporters. Several of them have been identified in *T. pseudonana* and shown to be located in the chloroplast or its bounding membranes (Moog et al. 2015). A search of our *T. oceanica* proteome showed one potential mitochondrial transporter, but so far, there is no experimental evidence that any of these is located in the mitochondrial outer membrane.

Interestingly, Hockin et al (2012) postulated that *T. pseudonana* increases glycolytic activity when nitrogen starved. However, when we mapped the involved proteins in *T. pseudonana* to their cellular target compartments, a regulation of isoenzymes similar to the response of Cu-limited *T. oceanica* emerged (i.e. PK down-regulated in mitochondria and up-regulated in the cytosol, Fig. S 3, Table S 4). Thus, the coordinated regulation of particular glycolytic isoenzymes to distribute NAD(P)H reducing equivalents and/or ATP production might be a general trait in diatoms.

### Nitrogen metabolism is essential for Fd^red^ oxidation

Another striking feature in the response to Cu limitation in *T. oceanica* was the up-regulation of nitrogen acquisition and assimilation as seen in the increased expression of 6 out of 8 proteins involved as well as the electron donor/acceptor ferredoxin (Fig. 5, Table S 6, overview Fig. S 5). In plants, nitrogen assimilation is an important sink for excess NAD(P)H (Hoefnagel et al., 1998). In *T. oceanica*, the increased expression may alleviate the stress incurred by low Cu, namely by re-oxidizing Fd^red^ in the chloroplast. This could be achieved via up-regulation of only those NiR isoenzymes that use Fd^red^ as their cofactor (To00016, To02363). Glutamine synthase (GSII, To31900) and the Fd^red^-dependent glutamate synthase (GOGAT, To13288) were also up-regulated, thereby potentially easing the chloroplast electron pressure. The importance of this proposed strategy for Cu-limited cells is highlighted by the fact that both the membrane-bound urea (To31656) and nitrate (To04919) transporters are among the 15 highest up-regulated proteins in our dataset.

### Counteracting reactive oxygen species – glutathione, thioredoxin, and superoxide dismutases

An enhanced nitrogen assimilation increases glutamate, which can be incorporated into (or be a precursor of) glutathione (GSH, γ-L-glutamyl-L-cysteine-glycine) to detoxify ROS via either direct scavenging or the ascorbate-glutathione cycle (Foyer and Noctor, 2011). Glutathione biosynthesis involves: (1) the cytosolic glutamate cysteine ligase (GCL, also known as γ-glutamylcysteine synthase, GCS) which combines glutamate and cysteine to γ-glutamyl-L-cysteine and (2) the plastid glutathione synthase (GSS) which adds glycine. Strikingly, both proteins were up-regulated in Cu-limited *T. oceanica*. However, in plants, the rate-limiting step in glutathione production is cysteine biosynthesis (Zechmann, 2014). Under Cu limitation, two chloroplast cysteine synthase isoenzymes were up-regulated (CS, To27524 and To10442; Fig. 7, Table 3, Table S 7) suggesting an increase in glutathione production. Furthermore, glutathione-S-transferase was one of the most highly up-regulated proteins (GST, To09062), which would be able to add glutathione to nucleophilic groups to detoxify oxidative stress (Gallogly and Mieyal, 2007). The up-regulation of glutathione reductase (GR, To07268), which oxidizes the over-abundant NAD(P)H in the chloroplast further supports our hypothesis that in *T. oceanica* glutathione counteracts ROS.

Thioredoxins (TXN) are important redox regulators in plants, especially in the chloroplast (Balmer et al., 2003), although their role in diatoms is unclear (Weber et al., 2009). In *T. oceanica*, the levels of three thioredoxins were increased, and each one was targeted to a different compartment: the chloroplast (TXN, To31425), the cytosol (To05213), and the mitochondria (To31425).

Another defence mechanism against ROS is the production of superoxide dismutases (SOD), which catalyze the conversion of superoxide radicals into hydrogen peroxide and oxygen. Of the three SODs identified in Cu-limited cultures, two were up-regulated: chloroplast Mn/Fe-SOD (To02860) and cytosolic Ni-SOD (To10112). Thus, under Cu limitation, cells may be able to control ROS levels by increasing the expression of both glutathione and SODs. The increase of thioredoxin isoenzymes in all three major cellular compartments (i.e. cytosol, chloroplast, mitochondria) points to their involvement in sensing the cellular redox state and regulating excess NAD(P)H.

### The malate shunt drains NAD(P)H reducing equivalents from the chloroplast to the mitochondria, thus integrating the nitrogen and carbon metabolisms

The efficiency of photosynthesis (both electron transport and carbon fixation) depends on an adequate supply of ATP/ADP and NAD(P)H/NAD(P)^+^ (Allen, 2002). In plants, the malate shunt can channel excess NAD(P)H reducing equivalents from the chloroplast to other cellular compartments, via the differential regulation of malate dehydrogenase (MDH) isoenzymes (Heineke et al., 1991; Scheibe, 2004). In this process, NAD(P)H in the chloroplast reduces oxaloacetate to malate, a compound that can be transported across membranes and re-oxidized, resulting in the production of NAD(P)H in the target compartment. NAD(P)H can then be used in reactions such as nitrate reduction in the cytosol or ATP production in the mitochondria.

In diatoms, the interaction between the chloroplast and mitochondria is expected to be multifaceted, possibly with direct exchange of ATP/ADP (Bailleul et al., 2015) and indirect exchange of NAD(P)H via the ornithine/glutamate shunt (Broddrick et al., 2019; Levering et al., 2016) and the malate/aspartate shunt (Bailleul et al., 2015; Prihoda et al., 2012). Some support for the spatial interconnectedness between chloroplast and mitochondria in diatoms has been reported recently (Flori et al., 2017). However, the location of the potential transporters needed (e.g. malate/2-oxoglutarate antiporter, glutamate/aspartate antiporter) have yet to be proven (Bailleul et al., 2015; Kim et al., 2016).

The proteomic patterns we present here support the existence and activation of the malate shunt in *T. oceanica* in response to low Cu. We observe the increased expression of chloroplast and mitochondrial MDH (MDH2, To30817; MDH1, To03405), as well as mitochondrial aspartate aminotransferase (AAT2, To15049, Fig. 6). As described by Kim et al (2016) in *P. tricornutum*, chloroplast oxaloacetate (OAA) is reduced to malate via MDH2. Malate is then transported into the mitochondria via a putative malate/2-oxoglutarate antiporter. NAD(P)H reducing equivalents are released in the mitochondria via the re-oxidation of malate to OAA by mitochondrial MDH1. In turn, mitochondrial AAT2 transfers an amine group from glutamate to OAA, thereby releasing aspartate and 2-oxoglutarate into the mitochondria. To close the cycle, aspartate is transported back, via a glutamate/aspartate antiporter, into the chloroplast where the plastidial AAT isoenzyme would resupply OAA (Kim et al., 2016). However, in *T. oceanica,* chloroplast AAT was significantly down-regulated. We suggest that chloroplast OAA, the substrate for MDH2, would be resupplied in the chloroplast via the ATP-dependent carboxylation of pyruvate due to the significant up-regulation of pyruvate carboxylase (PC). This would lead to a net decrease of NAD(P)H in the chloroplast and a net increase of NAD(P)H in the mitochondria. Furthermore, the channeling of NAD(P)H reducing equivalents towards respiration, instead of the Calvin-Benson-Bassham cycle, is supported by a 66% decreased in C fixation, while respiration rates remained constant (Hippmann et al, 2017).

The expected increase in 2-oxoglutarate and aspartate in the mitochondria, due to an up-regulation of mitochondrial AAT2, could be helpful for the cell. If the putative malate/2-oxoglutarate antiporter is indeed involved in the malate shunt, 2-oxoglutarate would be transported back into the chloroplast. As chloroplast AAT is down-regulated, 2-oxoglutarate could be used as a substrate for the up-regulated Fd-dependent glutamate synthase (GOGAT) in nitrogen assimilation. Any surplus 2-oxoglutarate in the mitochondria could feed into the citrate cycle. Fittingly, aconitase (To20545) and isocitrate dehydrogenase (To34595), the two proteins involved in the citrate cycle immediately before 2-oxoglutarate, were both significantly down-regulated (Fig. 3).

Mitochondrial aspartate can be channelled into the urea cycle, where it will produce argininosuccinate, which can then be diverted back into the mitochondrial citrate cycle as fumarate via the aspartate/fumarate shunt (Allen et al., 2011). Thus, even though two of the first three steps in the citrate cycle were down-regulated, we hypothesize that the malate shunt in combination with the urea cycle would ensure the continuation of this vital metabolic pathway by supplying it with essential carbon skeletons, i.e. 2-oxoglutarate and fumarate.

In addition to the malate shunt, other pathways have been proposed to alleviate electron pressure in diatoms. In *P. tricornutum*, modelling experiments suggest the prevalence of the glutamine-ornithine shunt over the malate shunt (Broddrick et al., 2019). However, none of the homologs involved in this shunt were identified in Cu-limited *T. oceanica* (e.g. n-acetyl-γ-glutamyl-phosphate reductase; n-acetylornithine aminotransferase). Furthermore, the activation of alternative oxidase (AOX) in Fe-limited *P. tricornutum* to alleviate electron stress in the impaired mitochondrial respiration (Allen et al., 2008) was not observed in Cu-limited *T. oceanica* (Hippmann et al., 2017). Future research is needed to elucidate the regulation of shuttle system/compartmental cross talks in diatoms.

### Conclusions

The success of diatoms in the modern ocean is thought to be due to their complex genomic makeup, and their successful integration and versatility of metabolic pathways. This was exemplified in the present study, where our proteomic data suggest how interaction among metabolic pathways act to maximize growth in *T. oceanica* (CCMP 1003) acclimated to severe Cu-limiting conditions. The differential expression of glycolytic isoenzymes located in the chloroplast and mitochondria may enable them to channel both excess electrons and/or ATP between these compartments. We found additional evidence for chloroplast-mitochondrial cross-talk in the reciprocal expression of chloroplast and mitochondrial isozymes involved in the proposed malate shunt, which could result in transferring both NAD(P)H reducing equivalents and carbon skeletons from the chloroplast to the mitochondria. The up-regulation of ferredoxin (Fdx) was correlated with up-regulation of plastidial Fdx-dependent isoenzymes involved in nitrogen assimilation as well as enzymes involved in glutathione synthesis, thus integrating nitrogen uptake and metabolism with photosynthesis and oxidative stress resistance.

## Methods

Cultures of the centric diatom *T. oceanica* strains CCMP 1003 and CCMP 1005 (here referred to as TO03 and TO05, respectively) were grown under Cu-replete and Cu-limited conditions. Proteins were extracted, purified, and analyzed by LC-MS/MS as detailed in Hippmann *et al.*(2017). The mass spectrometry proteomics data have been deposited to the ProteomeXchange Consortium via the PRIDE (Vizcaíno et al., 2016) partner repository with the dataset identifier PXD006237.

### Statistical analysis of differential protein expression

As described in Hippmann et al (2017), peptides were labelled with different isotopologues of formaldehyde depending on their nutrient regime (i.e. control, lowCu, lowFeCu), then mixed in a 1:1:1 ratio and analyzed by LC-MS/MS. Differential expression is then derived from the ratio of the intensities (area under the curve) of the light and heavy peaks for each peptide (see schematic in Fig. S 1). All three nutrient regimes (control, lowCu, lowFeCu) were processed together to be as consistent as possible and to decrease the number of false positives. However, in the present study we discuss the lowCu data only.

We defined two levels of statistically significant difference in expression: 1) greater than or equal to 2-fold (highly regulated), or 2) between 1.3- and 2-fold (regulated). In addition, the result must be found in at least two of the three biological replicates and result in a *p*-value of <0.05 for the z test, determining significant difference of the average ratios between treatments, taking the variance into account. Additionally, any protein that had a differential expression ratio of >10 (up-regulated) or <0.1 (down-regulated) in at least one biological replicate was considered to be an all-or-nothing response and was included in the ‘significantly changed’ set.

### Protein annotation and targeting prediction

Predicted proteins from both the publicly available genome of TO05 (CCMP1005) and our transcriptome of TO03 (CCMP 1003) were searched against a comprehensive sequence database, PhyloDB for functional annotation using BlastP. PhyloDB version 1.076 consists of 24,509,327 peptides from 19,962 viral, 230 archaeal, 4,910 bacterial, and 894 eukaryotic taxa. It includes peptides from the 410 taxa of the Marine Microbial Eukaryotic Transcriptome Sequencing Project (http://marinemicroeukaryotes.org/), as well as peptides from KEGG, GenBank, JGI, ENSEMBL, CAMERA, and various other repositories. To predict gene localization for proteins involved in carbon and nitrogen metabolism, four *in silico* strategies were followed: 1) sequences of candidate genes were compared to the publicly available chloroplast genomes of *T. oceanica* (CCMP 1005) (Lommer et al., 2010) and *T. pseudonana* (Armbrust et al., 2004; Oudot-Le Secq et al., 2007), 2) the diatom-specific chloroplast targeting sequence software ASAFind (Gruber et al., 2015) was used in conjunction with SignalP 3.0 (Petersen et al., 2011) to find nuclear encoded, chloroplast targeted proteins, 3) SignalP and TargetP (Emanuelsson et al., 2007) were used for mitochondrial targeting, and 4) comparison with curated subcellular locations of the closest homologs in *T. pseudonana*, *P. tricornutum*, and *Fragilariopsis cylindricus* genomes. We acknowledge that deducing cellular targeting via comparison to predicted or experimentally verified proteins in other diatoms can be challenging, as homologs can be found in different compartments depending on species (Gruber et al., 2015; Gruber and Kroth, 2017; Schober et al., 2019). The corresponding names of all protein abbreviations used throughout the present study (text and figures) are given in Table 1.

## Acknowledgements

We wish to thank Angele Arrieta and Michael Murphy (Department of Microbiology and Immunology, UBC) for critical reading of the manuscript. This work was financially supported by the Natural Sciences and Engineering Council of Canada Discovery Grants Program to M. T. Maldonado and L. J. Foster).

## Supplemental Tables and Figures

Table S 1: Comparison of cellular localization of various carbon metabolic pathways. (modified from (Gruber and Kroth, 2017).

Table S 2: Differential expression and predicted cellular location of proteins involved in the Calvin-Benson-Bassham cycle in TO03 and TO05 cultured in low Cu conditions *vs*. control.

Table S 3: Proteins with triose-phosphate transporter PFAM and their expression in TO03 and TO05 in response to Cu limitation *vs*. control

Table S 4: Differential expression and predicted cellular location of proteins involved in glycolysis in TO03 and TO05 cultured in low Cu conditions *vs*. control

Table S 5: Differential expression and predicted cellular location of proteins involved in the tricarboxylic acid (TCA) / citrate cycle in TO03 and TO05 cultured in low Cu conditions *vs*. control. (for diagram, see Fig. 3)

Table S 6: Differential expression and predicted cellular location of proteins involved in nitrogen metabolism including the urea cycle in TO03 and TO05 cultured in low Cu conditions *vs*. control.

Table S 7: Differential expression and predicted cellular location of proteins involved in glutathione metabolism in TO03 and TO05 cultured in low Cu conditions *vs*. control.

Table S 8: Differential expression and predicted cellular location of proteins involved in the putative malate shunt in TO03 and TO05 cultured in low Cu conditions *vs*. control

Table S 9: Differential expression and predicted cellular location of proteins involved in respiration in TO03 and TO05 cultured in low Cu conditions *vs*. control.

Fig. S 1: **Overview of Proteomic Method. A) Workflow, B) Table of preparation and mixing of samples** analyzed by LC-MS/MS.

Fig. S 2: **Clustal alignment of predicted amino acid sequences of TpMDH2** [Tp25953, original (TpMDH2_old) and EST extended (TpMDH2_new, Smith et al., 2012)] **and ToMDH2** (To30817). The new predicted cleavage sequence in TpMDH2_new is underlined.

Fig. S 3: **Comparison of differential expression of proteins involved in glycolysis under chronic Cu limitation and acute N limitation**.: A) *T. oceanica* (CCMP1003) under chronic Cu limitation (present study); B) *T. oceanica* (CCMP 1005) under chronic Cu limitation (present study, supplementary tables); C) *T. pseudonana* under acute N limitation (Hockin et al., 2012).

Fig. S 4: **Differential expression of proteins involved in pyruvate metabolism**.

Fig. S 5: **An overview of the proteomic response in the nitrogen and carbon metabolisms in *T. oceanica* (CCMP 1003) grown under Cu-limiting conditions**.

Fig. S 6: **An overview of the proteomic response in the nitrogen and carbon metabolisms in *T. oceanica* (CCMP 1005) grown under Cu-limiting conditions**.

Notes S 1: **Discussion on the contrasting adaptations to Cu limitation in the two strains of *T. oceanica* (CCMP1003 and CCMP1005).**

## Footnotes

1 AAH, AEA, LJF, BRG, and MTM planned and designed the research. AAH performed cell culturing, protein purification, statistical analysis, curated protein annotation, and targeting prediction. KMM performed protein purification and LC-MS/MS. JPM, under the guidance of AEA, created the RNAseq dataset and provided bioinformatic assistance to AAH. AAH, BRG, and MTM wrote the manuscript.

2 This research was funded by NSERC with grants to MTM and LJF.

3

## References

Allen, A.E., Dupont, C.L., Oborník, M., Horák, A., Nunes-Nesi, A., McCrow, J.P., Zheng, H., Johnson, D.A., Hu, H., Fernie, A.R., Bowler, C., 2011. Evolution and metabolic significance of the urea cycle in photosynthetic diatoms. Nature 473, 203–207. https://doi.org/10.1038/nature10074

Allen, A.E., LaRoche, J., Maheswari, U., Lommer, M., Schauer, N., Lopez, P.J., Finazzi, G., Fernie, A.R., Bowler, C., 2008. Whole-cell response of the pennate diatom Phaeodactylum tricornutum to iron starvation. Proceedings of the National Academy of Sciences 105, 10438–10443. https://doi.org/10.1073/pnas.0711370105

Allen, A.E., Moustafa, A., Montsant, A., Eckert, A., Kroth, P.G., Bowler, C., 2012. Evolution and Functional Diversification of Fructose Bisphosphate Aldolase Genes in Photosynthetic Marine Diatoms. Mol Biol Evol 29, 367–379. https://doi.org/10.1093/molbev/msr223

Allen, J.F., 2002. Photosynthesis of ATP—Electrons, Proton Pumps, Rotors, and Poise. Cell 110, 273–276. https://doi.org/10.1016/S0092-8674(02)00870-X

Annett, A.L., Lapi, S., Ruth, T.J., Maldonado, M.T., 2008. The effects of Cu and Fe availability on the growth and Cu:C ratios of marine diatoms. Limnol. Oceanogr. 53, 2451–2461. https://doi.org/10.4319/lo.2008.53.6.2451

Armbrust, E.V., 2009. The life of diatoms in the world’s oceans. Nature 459, 185–192. https://doi.org/10.1038/nature08057

Armbrust, E.V., Berges, J.A., Bowler, C., Green, B.R., Martinez, D., Putnam, N.H., Zhou, S., Allen, A.E., Apt, K.E., Bechner, M., Brzezinski, M.A., Chaal, B.K., Chiovitti, A., Davis, A.K., Demarest, M.S., Detter, J.C., Glavina, T., Goodstein, D., Hadi, M.Z., Hellsten, U., Hildebrand, M., Jenkins, B.D., Jurka, J., Kapitonov, V.V., Kröger, N., Lau, W.W.Y., Lane, T.W., Larimer, F.W., Lippmeier, J.C., Lucas, S., Medina, M., Montsant, A., Obornik, M., Parker, M.S., Palenik, B., Pazour, G.J., Richardson, P.M., Rynearson, T.A., Saito, M.A., Schwartz, D.C., Thamatrakoln, K., Valentin, K., Vardi, A., Wilkerson, F.P., Rokhsar, D.S., 2004. The Genome of the Diatom Thalassiosira Pseudonana: Ecology, Evolution, and Metabolism. Science 306, 79–86. https://doi.org/10.1126/science.1101156

Bailleul, B., Berne, N., Murik, O., Petroutsos, D., Prihoda, J., Tanaka, A., Villanova, V., Bligny, R., Flori, S., Falconet, D., Krieger-Liszkay, A., Santabarbara, S., Rappaport, F., Joliot, P., Tirichine, L., Falkowski, P.G., Cardol, P., Bowler, C., Finazzi, G., 2015. Energetic coupling between plastids and mitochondria drives CO2 assimilation in diatoms. Nature 524, 366–369. https://doi.org/10.1038/nature14599

Balmer, Y., Koller, A., Val, G. del, Manieri, W., Schürmann, P., Buchanan, B.B., 2003. Proteomics gives insight into the regulatory function of chloroplast thioredoxins. PNAS 100, 370–375. https://doi.org/10.1073/pnas.232703799

Bowler, C., Allen, A.E., Badger, J.H., Grimwood, J., Jabbari, K., Kuo, A., Maheswari, U., Martens, C., Maumus, F., Otillar, R.P., Rayko, E., Salamov, A., Vandepoele, K., Beszteri, B., Gruber, A., Heijde, M., Katinka, M., Mock, T., Valentin, K., Verret, F., Berges, J.A., Brownlee, C., Cadoret, J.-P., Chiovitti, A., Choi, C.J., Coesel, S., De Martino, A., Detter, J.C., Durkin, C., Falciatore, A., Fournet, J., Haruta, M., Huysman, M.J.J., Jenkins, B.D., Jiroutova, K., Jorgensen, R.E., Joubert, Y., Kaplan, A., Kröger, N., Kroth, P.G., La Roche, J., Lindquist, E., Lommer, M., Martin–Jézéquel, V., Lopez, P.J., Lucas, S., Mangogna, M., McGinnis, K., Medlin, L.K., Montsant, A., Secq, M.-P.O., Napoli, C., Obornik, M., Parker, M.S., Petit, J.-L., Porcel, B.M., Poulsen, N., Robison, M., Rychlewski, L., Rynearson, T.A., Schmutz, J., Shapiro, H., Siaut, M., Stanley, M., Sussman, M.R., Taylor, A.R., Vardi, A., von Dassow, P., Vyverman, W., Willis, A., Wyrwicz, L.S., Rokhsar, D.S., Weissenbach, J., Armbrust, E.V., Green, B.R., Van de Peer, Y., Grigoriev, I.V., 2008. The Phaeodactylum genome reveals the evolutionary history of diatom genomes. Nature 456, 239–244. https://doi.org/10.1038/nature07410

Broddrick, J.T., Du, N., Smith, S.R., Tsuji, Y., Jallet, D., Ware, M.A., Peers, G., Matsuda, Y., Dupont, C.L., Mitchell, B.G., Palsson, B.O., Allen, A.E., 2019. Cross-compartment metabolic coupling enables flexible photoprotective mechanisms in the diatom Phaeodactylum tricornutum. New Phytologist 222, 1364–1379. https://doi.org/10.1111/nph.15685

Emanuelsson, O., Brunak, S., von Heijne, G., Nielsen, H., 2007. Locating proteins in the cell using TargetP, SignalP and related tools. Nat. Protocols 2, 953–971. https://doi.org/10.1038/nprot.2007.131

Ewe, D., Tachibana, M., Kikutani, S., Gruber, A., Río Bártulos, C., Konert, G., Kaplan, A., Matsuda, Y., Kroth, P.G., 2018. The intracellular distribution of inorganic carbon fixing enzymes does not support the presence of a C4 pathway in the diatom Phaeodactylum tricornutum. Photosynth Res 137, 263–280. https://doi.org/10.1007/s11120-018-0500-5

Fabris, M., Matthijs, M., Rombauts, S., Vyverman, W., Goossens, A., Baart, G.J.E., 2012. The metabolic blueprint of Phaeodactylum tricornutum reveals a eukaryotic Entner– Doudoroff glycolytic pathway. The Plant Journal 70, 1004–1014. https://doi.org/10.1111/j.1365-313X.2012.04941.x

Field, C.B., Behrenfeld, M.J., Randerson, J.T., Falkowski, P., 1998. Primary Production of the Biosphere: Integrating Terrestrial and Oceanic Components. Science 281, 237–240. https://doi.org/10.1126/science.281.5374.237

Finazzi, G., Moreau, H., Bowler, C., 2010. Genomic insights into photosynthesis in eukaryotic phytoplankton. Trends in Plant Science 15, 565–572. https://doi.org/10.1016/j.tplants.2010.07.004

Flori, S., Jouneau, P.-H., Bailleul, B., Gallet, B., Estrozi, L.F., Moriscot, C., Bastien, O., Eicke, S., Schober, A., Bártulos, C.R., Maréchal, E., Kroth, P.G., Petroutsos, D., Zeeman, S., Breyton, C., Schoehn, G., Falconet, D., Finazzi, G., 2017. Plastid thylakoid architecture optimizes photosynthesis in diatoms. Nat Commun 8, 1–9. https://doi.org/10.1038/ncomms15885

Foyer, C.H., Noctor, G., 2011. Ascorbate and Glutathione: The Heart of the Redox Hub. Plant Physiol. 155, 2–18. https://doi.org/10.1104/pp.110.167569

Gallogly, M.M., Mieyal, J.J., 2007. Mechanisms of reversible protein glutathionylation in redox signaling and oxidative stress. Current Opinion in Pharmacology, Cancer/Immunomodulation 7, 381–391. https://doi.org/10.1016/j.coph.2007.06.003

Gruber, A., Kroth, P.G., 2017. Intracellular metabolic pathway distribution in diatoms and tools for genome-enabled experimental diatom research. Phil. Trans. R. Soc. B 372, 20160402. https://doi.org/10.1098/rstb.2016.0402

Gruber, A., Kroth, P.G., 2014. Deducing Intracellular Distributions of Metabolic Pathways from Genomic Data, in: Sriram, G. (Ed.), Plant Metabolism. Humana Press, Totowa, NJ, pp. 187–211. https://doi.org/10.1007/978-1-62703-661-0_12

Gruber, A., Rocap, G., Kroth, P.G., Armbrust, E.V., Mock, T., 2015. Plastid proteome prediction for diatoms and other algae with secondary plastids of the red lineage. Plant J 81, 519–528. https://doi.org/10.1111/tpj.12734

Gruber, A., Weber, T., Bártulos, C.R., Vugrinec, S., Kroth, P.G., 2009. Intracellular distribution of the reductive and oxidative pentose phosphate pathways in two diatoms. J. Basic Microbiol. 49, 58–72. https://doi.org/10.1002/jobm.200800339

Guo, J., Green, B.R., Maldonado, M.T., 2015. Sequence Analysis and Gene Expression of Potential Components of Copper Transport and Homeostasis in Thalassiosira pseudonana. Protist 166, 58–77. https://doi.org/10.1016/j.protis.2014.11.006

Guo, J., Lapi, S., Ruth, T.J., Maldonado, M.T., 2012. The Effects of Iron and Copper Availability on the Copper Stoichiometry of Marine Phytoplankton1. Journal of Phycology 48, 312–325. https://doi.org/10.1111/j.1529-8817.2012.01133.x

Heineke, D., Riens, B., Grosse, H., Hoferichter, P., Peter, U., Flügge, U.-I., Heldt, H.W., 1991. Redox Transfer across the Inner Chloroplast Envelope Membrane. Plant Physiol. 95, 1131–1137. https://doi.org/10.1104/pp.95.4.1131

Hippmann, A.A., Schuback, N., Moon, K.-M., McCrow, J.P., Allen, A.E., Foster, L.J., Green, B.R., Maldonado, M.T., 2017. Contrasting effects of copper limitation on the photosynthetic apparatus in two strains of the open ocean diatom Thalassiosira oceanica. PLOS ONE 12, e0181753. https://doi.org/10.1371/journal.pone.0181753

Hockin, N.L., Mock, T., Mulholland, F., Kopriva, S., Malin, G., 2012. The Response of Diatom Central Carbon Metabolism to Nitrogen Starvation Is Different from That of Green Algae and Higher Plants1[W]. Plant Physiol 158, 299–312. https://doi.org/10.1104/pp.111.184333

Hoefnagel, M.H.N., Atkin, O.K., Wiskich, J.T., 1998. Interdependence between chloroplasts and mitochondria in the light and the dark. Biochimica et Biophysica Acta (BBA) - Bioenergetics 1366, 235–255. https://doi.org/10.1016/S0005-2728(98)00126-1

Kim, J., Fabris, M., Baart, G., Kim, M.K., Goossens, A., Vyverman, W., Falkowski, P.G., Lun, D.S., 2016. Flux balance analysis of primary metabolism in the diatom Phaeodactylum tricornutum. Plant J 85, 161–176. https://doi.org/10.1111/tpj.13081

Kim, J.-W., Price, N.M., 2017. The influence of light on copper-limited growth of an oceanic diatom, Thalassiosira oceanica (Coscinodiscophyceae). J. Phycol. n/a-n/a. https://doi.org/10.1111/jpy.12563

Kong, L., M. Price, N., 2020. Identification of copper-regulated proteins in an oceanic diatom, Thalassiosira oceanica 1005. Metallomics 12, 1106–1117. https://doi.org/10.1039/D0MT00033G

Kroth, P.G., Chiovitti, A., Gruber, A., Martin-Jezequel, V., Mock, T., Parker, M.S., Stanley, M.S., Kaplan, A., Caron, L., Weber, T., Maheswari, U., Armbrust, E.V., Bowler, C., 2008. A Model for Carbohydrate Metabolism in the Diatom Phaeodactylum tricornutum Deduced from Comparative Whole Genome Analysis. PLOS ONE 3, e1426. https://doi.org/10.1371/journal.pone.0001426

Lelong, A., Bucciarelli, E., Hégaret, H., Soudant, P., 2013. Iron and copper limitations differently affect growth rates and photosynthetic and physiological parameters of the marine diatom Pseudo-nitzschia delicatissima. Limnol. Oceanogr. 58, 613–623. https://doi.org/10.4319/lo.2013.58.2.0613

Levering, J., Broddrick, J., Dupont, C.L., Peers, G., Beeri, K., Mayers, J., Gallina, A.A., Allen, A.E., Palsson, B.O., Zengler, K., 2016. Genome-Scale Model Reveals Metabolic Basis of Biomass Partitioning in a Model Diatom. PLOS ONE 11, e0155038. https://doi.org/10.1371/journal.pone.0155038

Lombardi, A.T., Maldonado, M.T., 2011. The effects of copper on the photosynthetic response of Phaeocystis cordata. Photosynth Res 108, 77–87. https://doi.org/10.1007/s11120-011-9655-z

Lommer, M., Roy, A.-S., Schilhabel, M., Schreiber, S., Rosenstiel, P., LaRoche, J., 2010. Recent transfer of an iron-regulated gene from the plastid to the nuclear genome in an oceanic diatom adapted to chronic iron limitation. BMC Genomics 11, 718. https://doi.org/10.1186/1471-2164-11-718

Maldonado, M.T., Allen, A.E., Chong, J.S., Lin, K., Leus, D., Karpenko, N., Harris, S.L., 2006. Copper-dependent iron transport in coastal and oceanic diatoms. Limnol. Oceanogr. 51, 1729–1743. https://doi.org/10.4319/lo.2006.51.4.1729

Maldonado, M.T., Hughes, M.P., Rue, E.L., Wells, M.L., 2002. The effect of Fe and Cu on growth and domoic acid production by Pseudo-nitzschia multiseries and Pseudo-nitzschia australis. Limnol. Oceanogr. 47, 515–526. https://doi.org/10.4319/lo.2002.47.2.0515

Moore, J.K., Doney, S.C., Lindsay, K., 2004. Upper ocean ecosystem dynamics and iron cycling in a global three-dimensional model. Global Biogeochem. Cycles 18, GB4028. https://doi.org/10.1029/2004GB002220

Moustafa, A., Beszteri, B., Maier, U.G., Bowler, C., Valentin, K., Bhattacharya, D., 2009. Genomic Footprints of a Cryptic Plastid Endosymbiosis in Diatoms. Science 324, 1724–1726. https://doi.org/10.1126/science.1172983

Mueller-Cajar, O., Stotz, M., Wendler, P., Hartl, F.U., Bracher, A., Hayer-Hartl, M., 2011. Structure and function of the AAA+ protein CbbX, a red-type Rubisco activase. Nature 479, 194–199. https://doi.org/10.1038/nature10568

Nelson, D.M., Tréguer, P., Brzezinski, M.A., Leynaert, A., Quéguiner, B., 1995. Production and dissolution of biogenic silica in the ocean: Revised global estimates, comparison with regional data and relationship to biogenic sedimentation. Global Biogeochem. Cycles 9, 359–372. https://doi.org/10.1029/95GB01070

Niyogi, K.K., 2000. Safety valves for photosynthesis. Current Opinion in Plant Biology 3, 455–460. https://doi.org/10.1016/S1369-5266(00)00113-8

Oborník, M., Green, B.R., 2005. Mosaic Origin of the Heme Biosynthesis Pathway in Photosynthetic Eukaryotes. Mol Biol Evol 22, 2343–2353. https://doi.org/10.1093/molbev/msi230

Oudot-Le Secq, M.-P.O.-L., Grimwood, J., Shapiro, H., Armbrust, E.V., Bowler, C., Green, B.R., 2007. Chloroplast genomes of the diatoms Phaeodactylum tricornutum and Thalassiosira pseudonana: comparison with other plastid genomes of the red lineage. Mol Genet Genomics 277, 427–439. https://doi.org/10.1007/s00438-006-0199-4

Peers, G., Price, N.M., 2006. Copper-containing plastocyanin used for electron transport by an oceanic diatom. Nature 441, 341–344. https://doi.org/10.1038/nature04630

Peers, G., Quesnel, S.-A., Price, N.M., 2005. Copper requirements for iron acquisition and growth of coastal and oceanic diatoms. Limnol. Oceanogr. 50, 1149–1158. https://doi.org/10.4319/lo.2005.50.4.1149

Petersen, T.N., Brunak, S., von Heijne, G., Nielsen, H., 2011. SignalP 4.0: discriminating signal peptides from transmembrane regions. Nat Meth 8, 785–786. https://doi.org/10.1038/nmeth.1701

Prihoda, J., Tanaka, A., Paula, W.B.M. de, Allen, J.F., Tirichine, L., Bowler, C., 2012. Chloroplast-mitochondria cross-talk in diatoms. J. Exp. Bot. 63, 1543–1557. https://doi.org/10.1093/jxb/err441

Río Bártulos, C., Rogers, M.B., Williams, T.A., Gentekaki, E., Brinkmann, H., Cerff, R., Liaud, M.-F., Hehl, A.B., Yarlett, N.R., Gruber, A., Kroth, P.G., van der Giezen, M., 2018. Mitochondrial Glycolysis in a Major Lineage of Eukaryotes. Genome Biol Evol 10, 2310–2325. https://doi.org/10.1093/gbe/evy164

Scheibe, R., 2004. Malate valves to balance cellular energy supply. Physiologia Plantarum 120, 21–26. https://doi.org/10.1111/j.0031-9317.2004.0222.x

Schober, A.F., Río Bártulos, C., Bischoff, A., Lepetit, B., Gruber, A., Kroth, P.G., 2019. Organelle Studies and Proteome Analyses of Mitochondria and Plastids Fractions from the Diatom Thalassiosira pseudonana. Plant Cell Physiol 60, 1811–1828. https://doi.org/10.1093/pcp/pcz097

Smith, S.R., Abbriano, R.M., Hildebrand, M., 2012. Comparative analysis of diatom genomes reveals substantial differences in the organization of carbon partitioning pathways. Algal Research 1, 2–16. https://doi.org/10.1016/j.algal.2012.04.003

Vizcaino, J.A., Csordas, A., del-Toro, N., Dianes, J.A., Griss, J., Lavidas, I., Mayer, G., Perez-Riverol, Y., Reisinger, F., Ternent, T., Xu, Q.-W., Wang, R., Hermjakob, H., 2016. 2016 update of the PRIDE database and its related tools. Nucleic Acids Res. 44, D447–456. https://doi.org/10.1093/nar/gkv1145

Weber, T., Gruber, A., Kroth, P.G., 2009. The Presence and Localization of Thioredoxins in Diatoms, Unicellular Algae of Secondary Endosymbiotic Origin. Molecular Plant 2, 468–477. https://doi.org/10.1093/mp/ssp010

Zechmann, B., 2014. Compartment-specific importance of glutathione during abiotic and biotic stress. Front Plant Sci 5. https://doi.org/10.3389/fpls.2014.00566

