## Supplemental File for "Proteomic analysis of metabolic pathways supports chloroplast-mitochondria cross-talk in a Cu-limited diatom"

Figure S 1: **Overview of Proteomic Method**. **A) Workflow: First the extracted proteins** are **trypsin digested. The resulting peptides are then** labelled via chemical dimethylation using light ($CH_2O$, for control treatment; green), medium ($CD_2O$, for low Cu treatment; blue), and heavy ($^{13}CD_2O$ for low FeCu treatment; red) isotopologues of formaldehyde as described previously (Boersema et al., 2009).After peptides of the respective treatments are labelled, they are mixed together in a 1:1:1 ratio and then analyzed together by LC-MS/MS. The differential expression of proteins is then derived from the ratio of the intensities (area under the curve, here depicted as the height of bars) of the light (green), medium (blue) and heavy (red) peaks for each peptide.

Figure S 2: **Clustal alignment of predicted amino acid sequences of TpMDH2** [Tp25953, original (TpMDH2_old) and EST extended (TpMDH2_new, Smith et al., 2012)] **and ToMDH2** (To30817). The new predicted cleavage sequence in TpMDH2_new is underlined

Figure S 3: **Comparison of differential expression of proteins involved in glycolysis under chronic Cu limitation and acute N limitation**.: A) *T. oceanica* (CCMP1003) under chronic Cu limitation (present study); B) *T. oceanica* (CCMP 1005) under chronic Cu limitation (present study, supplementary tables); C) *T. pseudonana* under acute N limitation (Hockin et al., 2012). Boxes indicate proteins with their abbreviated name (for full name, see **Error! Reference source not found.**). Dark red boxes indicate highly up-regulated proteins (TO03: > 2-fold, $p<0.05$; Tp: as per Hockin et al. 2012), light red boxes indicate 1.3- to 2-fold up-regulated proteins ($p<0.05$), white boxes indicate no significant regulation, dark blue boxes indicate highly down-regulated proteins (TO03: > 2-fold, $p<0.05$; Tp: as per Hockin et al. 2012), light blue boxes indicate 1.3 to 2-fold down-regulated proteins ($p<0.05$).

Figure S 4: **Differential expression of proteins involved in pyruvate metabolism**. Boxes indicate proteins with their abbreviated name and known *T. pseudonana* (Tp) and *T. oceanica* (To) homologs. The colours of the boxes indicate expression in TO03 under low Cu: dark red, highly up-regulated (>2-fold, $p<0.05$); light pink, up-regulated by 1.3 to 2-fold ($p<0.05$); dark blue, highly down-regulated (>2-fold, $p<0.05$); light blue, down-regulated by 1.3 to 2-fold ($p<0.05$); white, expressed in TO03; grey border around box, found in *T. oceanica* genome but not expressed in TO03 proteomic data; grey, dashed border around box, no putative homologs in the *T. oceanica* genome. [a]Tp and To model are not homologs

Figure S 5: **An overview of the proteomic response in the nitrogen and carbon metabolisms in *T. oceanica* (CCMP 1003) grown under Cu-limiting conditions.** Boxes indicate proteins with their abbreviated name. The colors of the boxes indicate expression in TO03 under low Cu: dark red, highly up-regulated (> 2-fold, $p<0.05$); light pink, up-regulated by 1.3 to 2-fold ($p<0.05$); dark blue, highly down-regulated (> 2-fold, $p<0.05$); light blue, down-regulated by 1.3 to 2-fold ($p<0.05$); white, expressed in TO03; grey border around box, found in *T. oceanica* genome but not detected in TO03 proteomic data; grey, dashed border around box, no putative homologs in the *T. oceanica* genome.

Figure S 6: **An overview of the proteomic response in the nitrogen and carbon metabolisms in *T. oceanica* (CCMP 1005) grown under Cu-limiting conditions.** Boxes indicate proteins with their abbreviated name. The colors of the boxes indicate expression in TO05 under low Cu: dark red, highly up-regulated (> 2-fold, $p<0.05$); light pink, up-regulated by 1.3 to 2-fold ($p<0.05$); dark blue, highly down-regulated (> 2-fold, $p<0.05$); light blue, down-regulated by 1.3

to 2-fold ($p<0.05$); white, expressed in TO05; grey border around box, found in *T. oceanica* genome but not detected in TO05 proteomic data; grey, dashed border around box, no putative homologs in the *T. oceanica* genome.

Table S 1: **Comparison of cellular localization of various carbon metabolic pathways.** (modified from (Gruber & Kroth, 2017)Table S 1: **Comparison of cellular localization of various carbon metabolic pathways.** (modified from (Gruber & Kroth, 2017)

Table S 2: **Differential expression and predicted cellular location of proteins involved in the Calvin-Benson-Bassham cycle** in TO03 and TO05 cultured in low Cu conditions *vs.* control.

Table S 3: **Proteins with triose-phosphate transporter PFAM and their expression in TO03 and TO05** in response to Cu limitation *vs.* control.

Table S 4: **Differential expression and predicted cellular location of proteins involved in glycolysis** in TO03 and TO05 cultured in low Cu conditions *vs.* control.

Table S 5: **Differential expression and predicted cellular location of proteins involved in the tricarboxylic acid (TCA) / citrate cycle** in TO03 and TO05 cultured in low Cu conditions *vs.*control. (for diagram, see Fig. 3, main text)

Table S 6: **Differential expression and predicted cellular location of proteins involved in nitrogen metabolism including the urea cycle** in TO03 and TO05 cultured in low Cu conditions *vs.* control.

Table S 7: **Differential expression and predicted cellular location of proteins involved in glutathione metabolism** in TO03 and TO05 cultured in low Cu conditions *vs.* control.

Table S 8: **Differential expression and predicted cellular location of proteins involved in the putative malate shunt** in TO03 and TO05 cultured in low Cu conditions *vs.* control.

Table S 9: **Differential expression and predicted cellular location of proteins involved in respiration** in TO03 and TO05 cultured in low Cu conditions *vs.* control.

**Notes S 1: Discussion on the contrasting adaptations to Cu limitation in the two strains of *T. oceanica* (CCMP1003 and CCMP1005).**

A

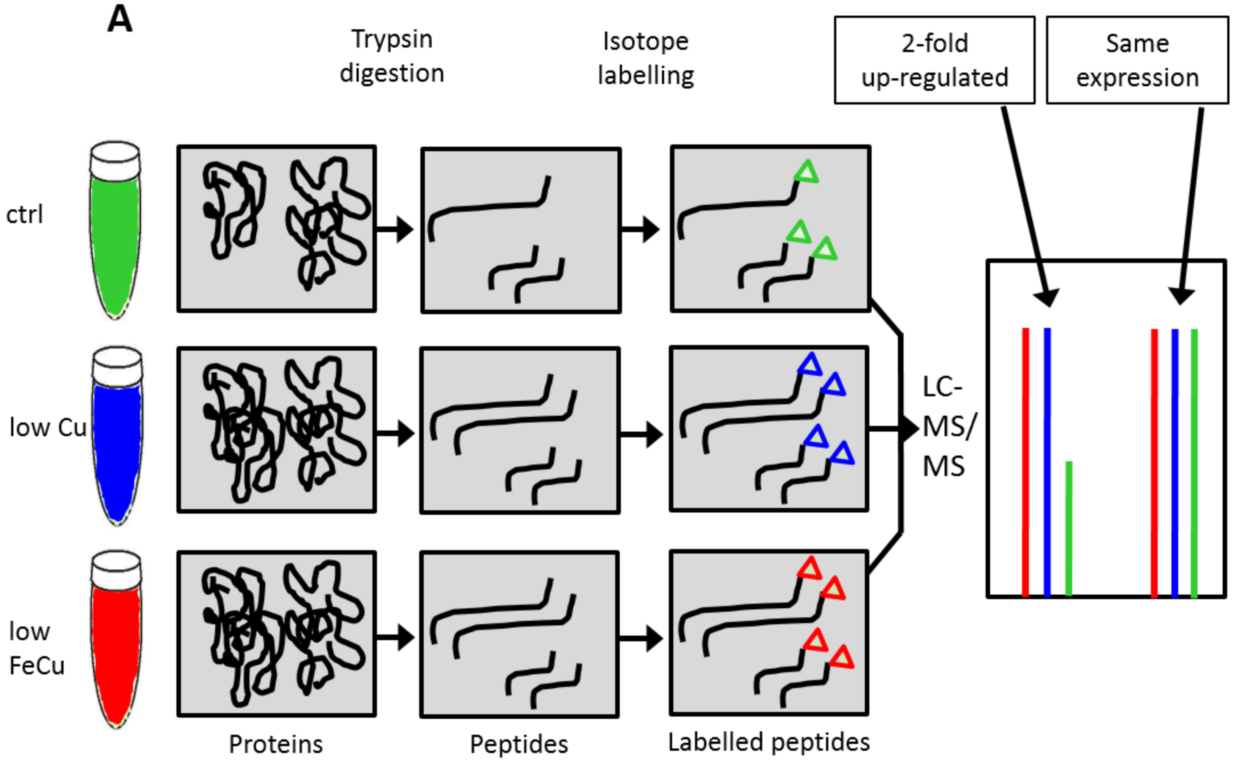


B

| Treatment (label) | Biological Replicate | | |
|---|---|---|---|
| Control (low) | 1 | 2 | 3 |
| lowCu (medium) | 1 | 2 | 3 |
| lowFeCu (heavy) | 1 | 2 | 3 |
| | ↓ | ↓ | ↓ |
| 1:1:1 mix of labelled samples | 1 | 2 | 3 |

Figure S 1: **Overview of Proteomic Method**. **A) Workflow**: First the extracted proteins are trypsin digested. The resulting peptides are then labelled via chemical dimethylation using light ($CH_2O$, for control treatment; green), medium ($CD_2O$, for low Cu treatment; blue), and heavy ($^{13}CD_2O$ for low FeCu treatment; red) isotopologues of formaldehyde as described previously (Boersema et al., 2009).After peptides of the respective treatments are labelled, they are mixed together in a 1:1:1 ratio and then analyzed together by LC-MS/MS. The differential expression of proteins is then derived from the ratio of the intensities (area under the curve, here depicted as the height of bars) of the light (green), medium (blue) and heavy (red) peaks for each peptide., **B) Table of preparation and mixing of samples** analyzed by LC-MS/MS. Each biological replicate (three per treatment) is labelled individually. Then one labelled sample of each treatment is mixed together in a 1:1:1 ratio, resulting in three separate biological replicate mixes to be analyzed by LC-MS/MS. Each of these three biological replicate mixes was then analyzed in technical duplicates (TO03) or triplicates (TO05).

```
CLUSTAL format alignment by MAFFT (v7.408)


TpMDH2_new      MLRKQAAAALTLTLLTPHLCIADAFLPPQSSVATFTHNHHNSKLENSSMSAASDLRAELL
TpMDH2_old      ------------------------------------------------------------
ToMDH2          ------------------------------------------------MSSSAEIRGELL


TpMDH2_new      ELIGKNPEGHKDATVRSHFASKLEAYSQAEGADPSDVIFAIGQTVFGVDL-PPPAKKSKT
TpMDH2_old      ------------------------------------------------------------
ToMDH2          KLIGEN--TNPDETVKSAFATKLKEYA-AAGGESNDVLFAIGKTVYGAELDQPPAKKAKA


TpMDH2_new      GAEDAPLPIVDFDYMKAFMKEVFLSYGVTPERAEVCSDVLIESDKRGIDSHGLGRLKPIY
TpMDH2_old      --------------MKAFMKEVFLSYGVTPERAEVCSDVLIESDKRGIDSHGLGRLKPIY
ToMDH2          APADE-LPVVDFDYMKAFMKDVFLSYGVTPENAEVCADVLIESDKRGIDSHGLGRLKPIY
                              ******:**********.****:***********************

TpMDH2_new      CDRMDDGILFPDKPIDIISESDTTALVDGNLGLGLYIGPHCMQMAIDKAKKHGVGFVAVR
TpMDH2_old      CDRMDDGILFPDKPIDIISESDTTALVDGNLGLGLYIGPHCMQMAIDKAKKHGVGFVAVR
ToMDH2          CDRMDNGILFPDKPIDIVSESETTALVDGNLGLGLYIGPHCMQMAIDKAKKHGVGFVAVR
                *****:***********:***:**************************************

TpMDH2_new      NSTHYGIAGYYATMATQQGCIGLTGTNARPSIAPTFGVEPMMGTNPLTFGIPSSDDFPFV
TpMDH2_old      NSTHYGIAGYYATMATQQGCIGLTGTNARPSIAPTFGVEPMMGTNPLTFGIPSSDDFPFV
ToMDH2          NSTHYGIAGYYATMASSQGCVGLTGTNARPSIAPTFGVEPMLGTNPLTFGIPSTDEWPFV
                ***************:.***:********************:***********:*::***

TpMDH2_new      IDCATSVNQRGKIEKYAREGVDTPRGAVIDDQGIERTDTDGILRDMVLGKCALTPVGGAG
TpMDH2_old      IDCATSVNQRGKIEKYAREGVDTPRGAVIDDQGIERTDTDGILRDMVLGKCALTPVGGAG
ToMDH2          IDCATSVNQRGKIERYAREGLDTPRGAVIDDQGIERTDTEGILRDMVLGKCALTPVGGAG
                **************:*****:*****************:*********************

TpMDH2_new      DKMGGYKGYGWATTVELLCTALQSGPWGEDICGVDRATGKPKPMPLGHFFLAIDIEKICP
TpMDH2_old      DKMGGYKGYGWATTVELLCTALQSGPWGEDICGVDRATGKPKPMPLGHFFLAIDIEKICP
ToMDH2          DKMGGYKGYGWATTVELLCTAFQSGPWGEDICGLDRATGKPKPMPLGHFFLAIDIEKLCP
                *********************:***********:***********************:**

TpMDH2_new      VDTFKKNSGEFLQALRDSKKAPNGPGRIWTAGEIENDARVERTAQGGMKVPIPLQKNMKA
TpMDH2_old      VDTFKKNSGEFLQALRDSKKAPNGPGRIWTAGEIENDARVERTAQGGMKVPIPLQKNMKA
ToMDH2          LDTFKKNSGEFLKALRESRKAPNGPGRIWTAGEPENDARVQRTEQGGMKVNVPLQKNMAA
                :***********:***:*:************** ******:** ****** :****** *

TpMDH2_new      LRDTRPGLKEKYVKLLFE-
TpMDH2_old      LRDTRPGLKEKYVKLLFE-
ToMDH2          LRESRPGLKEKYAKFPFES
                **::********.*: **
```

Figure S 2: **Clustal alignment of predicted amino acid sequences of TpMDH2** [Tp25953, original (TpMDH2_old) and EST extended (TpMDH2_new, Smith et al., 2012)] **and ToMDH2** (To30817). The new predicted cleavage sequence in TpMDH2_new is underlined

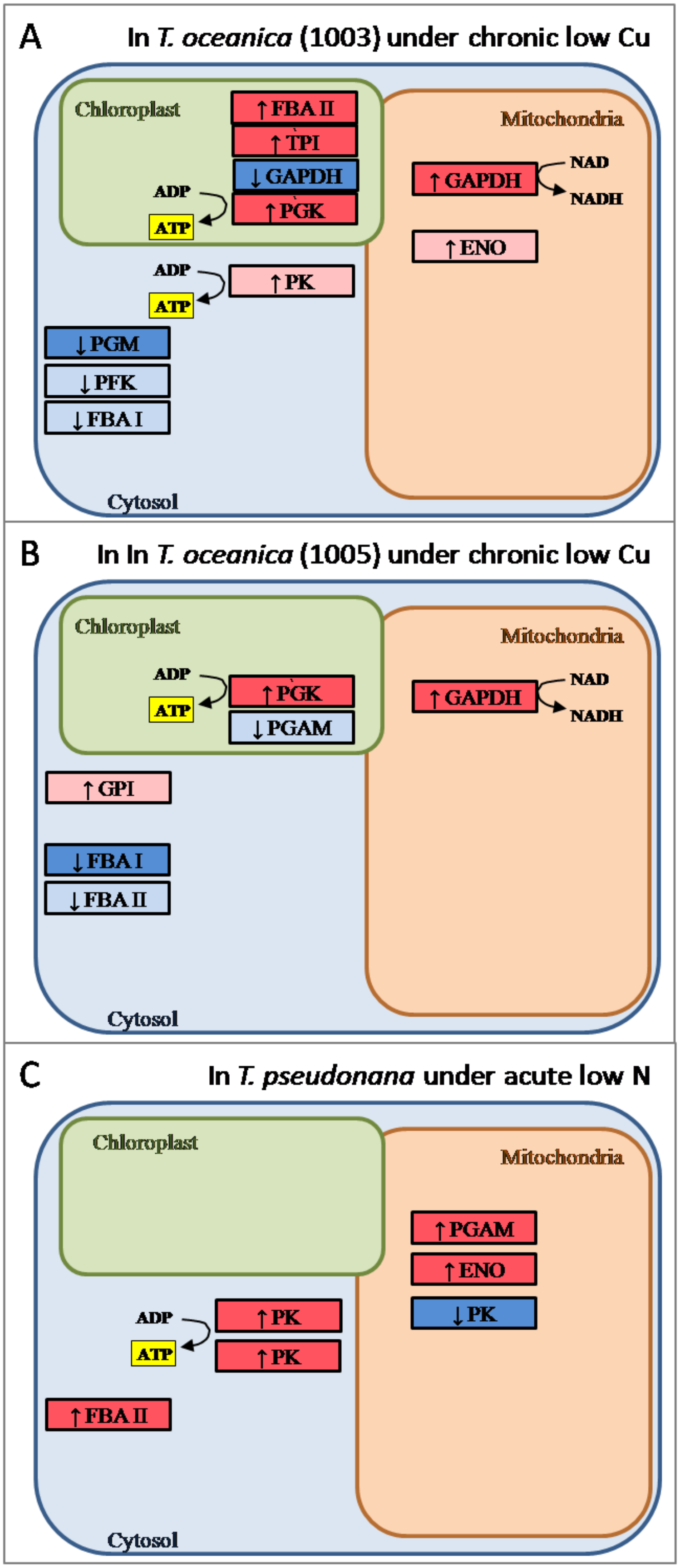
A
In T. oceanica (1003) under chronic low Cu
Chloroplast
↑FBA II
↑TPI
↓GAPDH
↑PGK
ADP
ATP
Mitochondria
↑GAPDH
NAD
NADH
↑ENO
ADP
ATP
↑PK
↓PGM
↓PFK
↓FBA I
Cytosol
B
In In T. oceanica (1005) under chronic low Cu
Chloroplast
ADP
ATP
↑PGK
↓PGAM
Mitochondria
↑GAPDH
NAD
NADH
↑GPI
↓FBA I
↓FBA II
Cytosol
C
In T. pseudonana under acute low N
Chloroplast
Mitochondria
↑PGAM
↑ENO
↓PK
ADP
ATP
↑PK
↑PK
↑FBA II
Cytosol

Figure S 3: **Comparison of differential expression of proteins involved in glycolysis under chronic Cu limitation and acute N limitation**.: A) *T. oceanica* (CCMP1003) under chronic Cu limitation (present study); B) *T. oceanica* (CCMP 1005) under chronic Cu limitation (present study, supplementary tables); C) *T. pseudonana* under acute N limitation (Hockin et al., 2012). Boxes indicate proteins with their abbreviated name (for full name, see **Error! Reference source not found.**). Dark red boxes indicate highly up-regulated proteins (TO03: > 2-fold, $p<0.05$; Tp: as per Hockin et al. 2012), light red boxes indicate 1.3- to 2-fold up-regulated proteins ($p<0.05$), white boxes indicate no significant regulation, dark blue boxes indicate highly down-regulated proteins (TO03: > 2-fold, $p<0.05$; Tp: as per Hockin et al. 2012), light blue boxes indicate 1.3 to 2-fold down-regulated proteins ($p<0.05$).

**Abbreviations:** EDA, 2-keto-3-deoxy phosphogluconate aldolase; EDD, 6-phosphogluconate dehydratase; ENO, enolase; F2BP, fructose-1-6-bisphosphatase; FBA I, fructose-bisphosphate aldolase class-I; FBA II, fructose-bisphosphate aldolase class-II; GAPDH, glyceraldehyde 3-phosphate dehydrogenase; GPI, glucose-6-phosphate isomerase; PFK, phosphofructokinase; PGAM, phosphoglycerate mutase; PGK, phosphoglycerate kinase; PGM, phosphoglucomutase; PK, pyruvate kinase; TPI, triose-phosphate isomerase

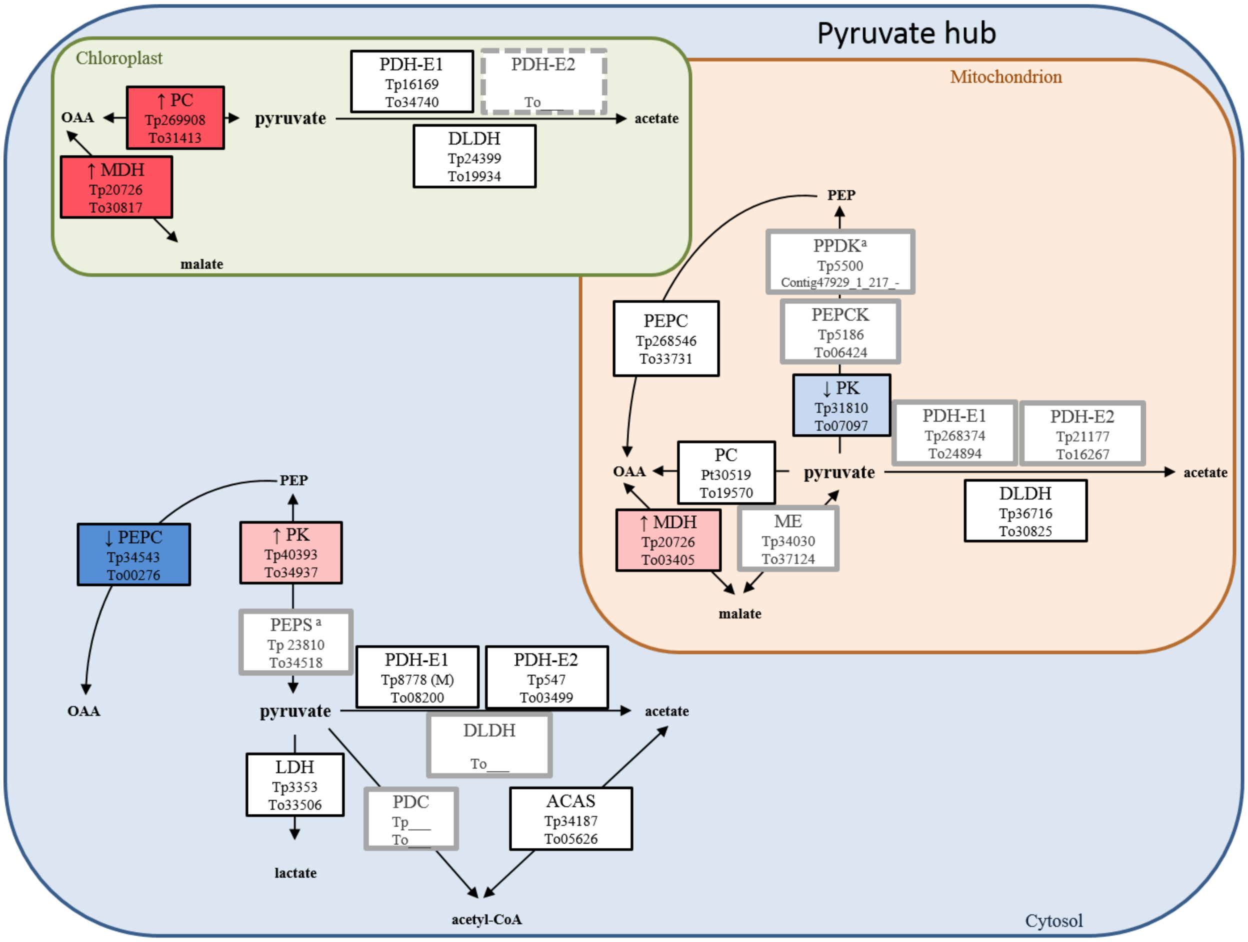


Figure S 4: **Differential expression of proteins involved in pyruvate metabolism**. Boxes indicate proteins with their abbreviated name and known *T. pseudonana* (Tp) and *T. oceanica* (To) homologs. The colours of the boxes indicate expression in TO03 under low Cu: dark red, highly up-regulated (>2-fold, $p<0.05$); light pink, up-regulated by 1.3 to 2-fold ($p<0.05$); dark blue, highly down-regulated (>2-fold, $p<0.05$); light blue, down-regulated by 1.3 to 2-fold ($p<0.05$); white, expressed in TO03; grey border around box, found in *T. oceanica* genome but not expressed in TO03 proteomic data; grey, dashed border around box, no putative homologs in the *T. oceanica* genome. [a]Tp and To model are not homologs

**Abbreviations:** ACAS, acetyl-CoA synthase; ACC, acetyl-CoA carboxylase; DLDH, dihydrolipoamide dehydrogenase; LDH, L-lactate dehydrogenase; MDH, malate dehydrogenase; ME, malic enzyme; PC, pyruvate carboxylase; PDH-E1, pyruvate dehydrogenase-E1 component; PDH-E2, pyruvate dehydrogenase E2 component (dihydrolipoamide acetyltransferase); PEPC, phosphoenolpyruvate carboxylase; PEPCK, phosphoenolpyruvate carboxykinase; PEPS, phosphoenolpyruvate synthase; PK, pyruvate kinase; PPDK, pyruvate, phosphate dikinase

TO03

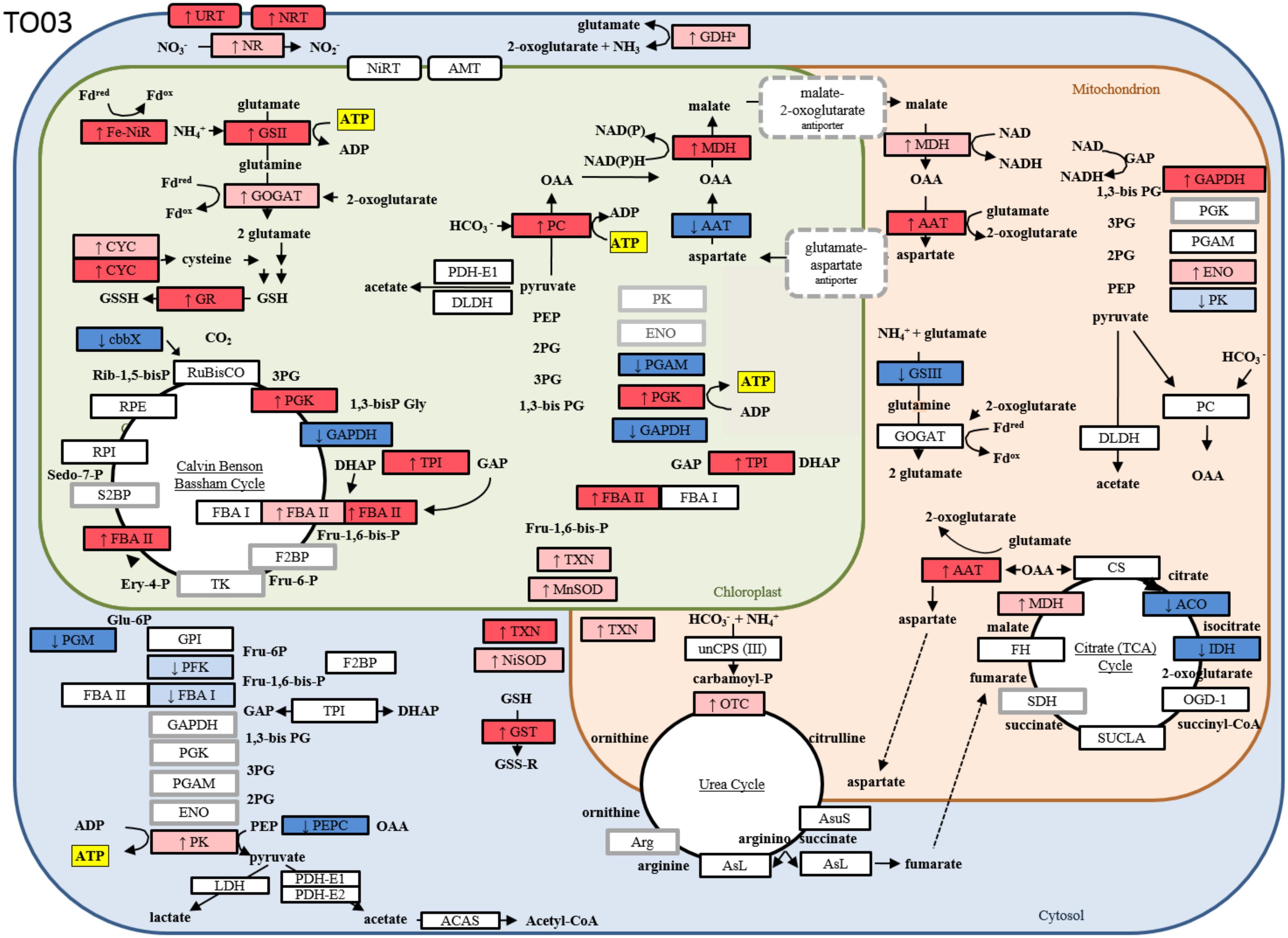
↑ URT
↑ NRT
$NO_3^-$
↑ NR
$NO_2^-$
glutamate
2-oxoglutarate + $NH_3$
↑ GDH[a]
NiRT
AMT
Mitochondrion
Fd[red]
Fd[ox]
↑ Fe-NiR
$NH_4^+$
glutamate
↑ GSII
ATP
ADP
glutamine
Fd[red]
Fd[ox]
↑ GOGAT
2-oxoglutarate
2 glutamate
↑ CYC
↑ CYC
cysteine
GSSH
↑ GR
GSH
↓ cbbX
$CO_2$
Rib-1,5-bisP
RuBisCO
3PG
↑ PGK
1,3-bisP Gly
RPE
↓ GAPDH
RPI
Sedo-7-P
Calvin Benson Bassham Cycle
DHAP
↑ TPI
GAP
S2BP
FBA I
↑ FBA II
↑ FBA II
Fru-1,6-bis-P
↑ FBA II
F2BP
Ery-4-P
TK
Fru-6-P
malate
NAD(P)
NAD(P)H
↑ MDH
OAA
OAA
$HCO_3^-$
↑ PC
ADP
ATP
↓ AAT
aspartate
PDH-E1
acetate
DLDH
pyruvate
PEP
2PG
3PG
1,3-bis PG
PK
ENO
↓ PGAM
↑ PGK
ATP
ADP
↓ GAPDH
GAP
↑ TPI
DHAP
↑ FBA II
FBA I
Fru-1,6-bis-P
↑ TXN
↑ MnSOD
Chloroplast
malate-
2-oxoglutarate
antiporter
malate
glutamate-
aspartate
antiporter
↑ MDH
NAD
NADH
OAA
↑ AAT
glutamate
2-oxoglutarate
aspartate
NAD
GAP
NADH
1,3-bis PG
↑ GAPDH
PGK
3PG
PGAM
2PG
↑ ENO
PEP
↓ PK
pyruvate
$NH_4^+$ + glutamate
↓ GSIII
glutamine
2-oxoglutarate
Fd[red]
GOGAT
Fd[ox]
2 glutamate
$HCO_3^-$
PC
DLDH
acetate
OAA
2-oxoglutarate
glutamate
↑ AAT
OAA
CS
citrate
aspartate
↑ MDH
↓ ACO
malate
isocitrate
FH
Citrate (TCA) Cycle
↓ IDH
fumarate
2-oxoglutarate
SDH
OGD-1
succinate
SUCLA
succinyl-CoA
Glu-6P
↓ PGM
GPI
Fru-6P
↓ PFK
F2BP
Fru-1,6-bis-P
FBA II
↓ FBA I
GAP
TPI
DHAP
GAPDH
1,3-bis PG
PGK
3PG
PGAM
2PG
ENO
ADP
PEP
↓ PEPC
OAA
ATP
↑ PK
pyruvate
LDH
PDH-E1
PDH-E2
lactate
acetate
ACAS
Acetyl-CoA
↑ TXN
↑ NiSOD
GSH
↑ GST
GSS-R
↑ TXN
$HCO_3^-$ + $NH_4^+$
unCPS (III)
carbamoyl-P
↑ OTC
ornithine
citrulline
Urea Cycle
aspartate
ornithine
AsuS
Arg
arginino succinate
arginine
AsL
AsL
fumarate
Cytosol

Figure S 5: **An overview of the proteomic response in the nitrogen and carbon metabolisms in *T. oceanica* (CCMP 1003) grown under Cu-limiting conditions.** Boxes indicate proteins with their abbreviated name. The colors of the boxes indicate expression in TO03 under low Cu: dark red, highly up-regulated (> 2-fold, $p<0.05$); light pink, up-regulated by 1.3 to 2-fold ($p<0.05$); dark blue, highly down-regulated (> 2-fold, $p<0.05$); light blue, down-regulated by 1.3 to 2-fold ($p<0.05$); white, expressed in TO03; grey border around box, found in *T. oceanica* genome but not detected in TO03 proteomic data; grey, dashed border around box, no putative homologs in the *T. oceanica* genome.

**Abbreviations:** AAT, aspartate aminotransferase; ACO, aconitase hydratase; Arg, Arginase; AsL, argininosuccinate lyase; AsuS, argininosuccinate synthase; cbbX, rubisco expression protein; CS, citrate synthase; DLDH, dihydrolipoamide dehydrogenase; EDA, 2-keto-3-deoxy phosphogluconate aldolase; EDD, 6-phosphogluconate dehydratase; ENO, Enolase; F2BP, Fructose-1-6-bisphosphatase; FBA I, fructose-bisphosphate aldolase class-I; FBA II, fructose-bisphosphate aldolase class-II; Fe-NiR, nitrite reductase (ferredoxin-dependent); FH, fumarate hydratase; GAPDH, glyceraldehyde 3-phosphate dehydrogenase; GDH, glutamate dehydrogenase; GOGAT, glutamate synthase; GPI, glucose-6-phosphate isomerase; GSII, glutamine synthetase; GSIII, glutamine synthetase; IDH, isocitrate dehydrogenase; MDH, malate dehydrogenase; NR, nitrate reductase; NRT, nitrate/nitrite transporter; OGD, 2-oxoglutarate dehydrogenase; OTC, ornithine carbamoyltransferase; PC, pyruvate carboxylase; PEPC, phosphoenolpyruvate carboxylase; PFK, phosphofructokinase; PGAM, phosphoglycerate mutase; PGK, phosphoglycerate kinase; PGM, phosphoglucomutase; PK, pyruvate kinase; RPE, ribulose-5-phosphate epimerase; RPI, ribose-5-Phosphate-isomerase; RuBisCO, ribulose-bisphosphate carboxylase; SUCLA, Succinate CoA synthetase; TPI, triose-phosphate isomerase; TPI, triose-phosphate isomerase; unCPS (CPSase III), carbamoyl-phosphate synthase; URT, Na/urea-polyamine transporter.

TO05

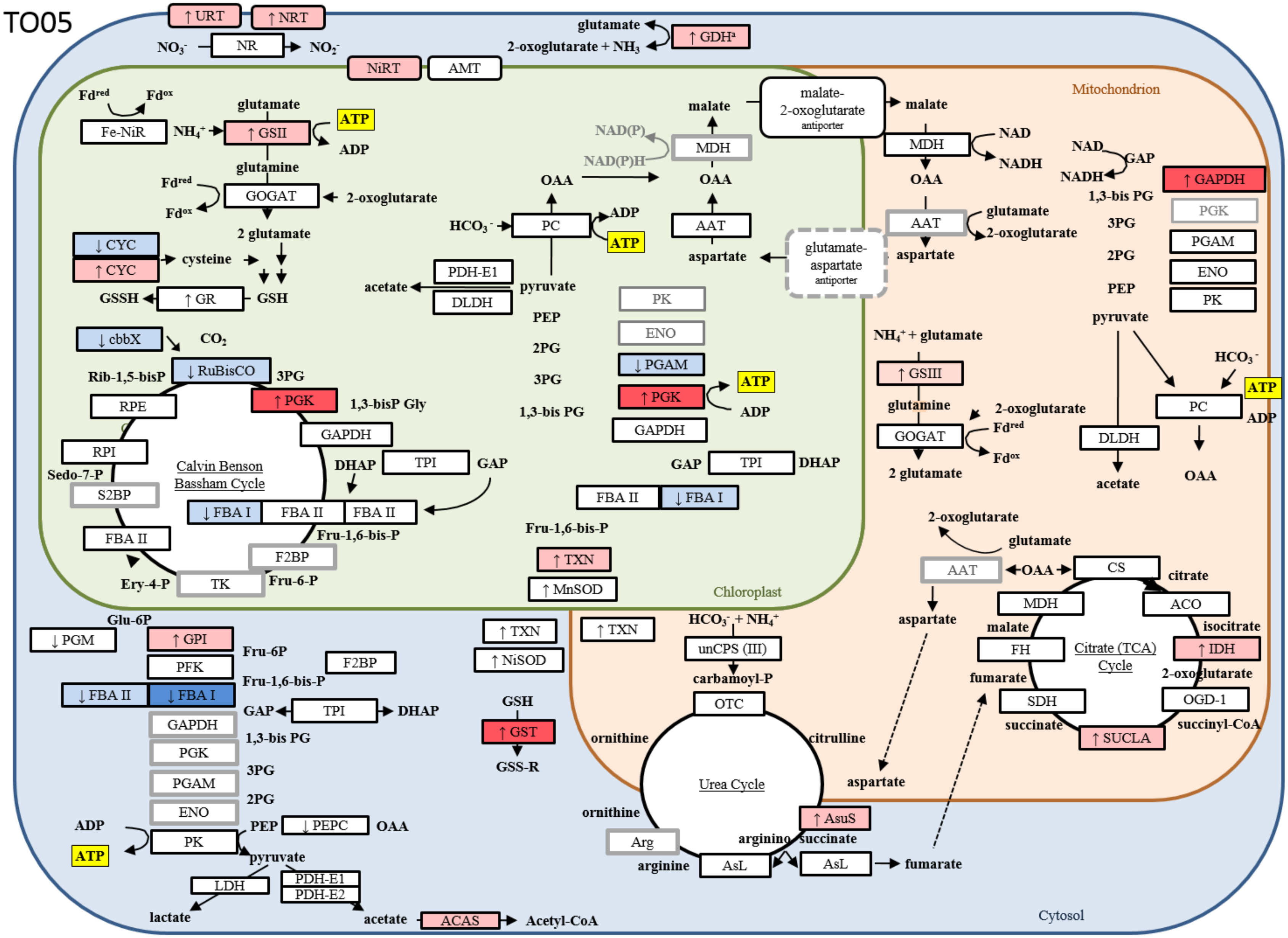
↑ URT
↑ NRT
$NO_3^-$
NR
$NO_2^-$
glutamate
2-oxoglutarate + $NH_3$
↑ GDH[a]
NiRT
AMT
Mitochondrion
Chloroplast
Cytosol
$Fd^{red}$
$Fd^{ox}$
Fe-NiR
$NH_4^+$
glutamate
↑ GSII
ATP
ADP
glutamine
GOGAT
2-oxoglutarate
2 glutamate
↓ CYC
↑ CYC
cysteine
GSSH
↑ GR
GSH
↓ cbbX
$CO_2$
Rib-1,5-bisP
↓ RuBisCO
3PG
↑ PGK
1,3-bisP Gly
RPE
GAPDH
RPI
Calvin Benson Bassham Cycle
DHAP
TPI
GAP
Sedo-7-P
S2BP
↓ FBA I
FBA II
FBA II
FBA II
Fru-1,6-bis-P
F2BP
Ery-4-P
TK
Fru-6-P
$HCO_3^-$
PC
ADP
ATP
OAA
PDH-E1
DLDH
acetate
pyruvate
PEP
2PG
3PG
1,3-bis PG
Fru-1,6-bis-P
↑ TXN
↑ MnSOD
NAD(P)
NAD(P)H
MDH
malate
OAA
AAT
aspartate
malate-2-oxoglutarate antiporter
glutamate-aspartate antiporter
PK
ENO
↓ PGAM
↑ PGK
ATP
ADP
GAPDH
GAP
TPI
DHAP
FBA II
↓ FBA I
malate
MDH
NAD
NADH
OAA
AAT
glutamate
2-oxoglutarate
aspartate
NAD
GAP
NADH
1,3-bis PG
↑ GAPDH
PGK
3PG
PGAM
2PG
ENO
PEP
PK
pyruvate
$NH_4^+$ + glutamate
↑ GSIII
glutamine
2-oxoglutarate
$Fd^{red}$
GOGAT
$Fd^{ox}$
2 glutamate
$HCO_3^-$
ATP
PC
ADP
DLDH
acetate
OAA
2-oxoglutarate
glutamate
AAT
OAA
CS
citrate
aspartate
MDH
ACO
malate
isocitrate
FH
Citrate (TCA) Cycle
↑ IDH
fumarate
2-oxoglutarate
SDH
OGD-1
succinate
↑ SUCLA
succinyl-CoA
Glu-6P
↓ PGM
↑ GPI
Fru-6P
PFK
F2BP
Fru-1,6-bis-P
↓ FBA II
↓ FBA I
GAP
TPI
DHAP
GAPDH
1,3-bis PG
PGK
3PG
PGAM
2PG
ENO
ADP
PEP
↓ PEPC
OAA
ATP
PK
pyruvate
LDH
PDH-E1
PDH-E2
lactate
acetate
ACAS
Acetyl-CoA
↑ TXN
↑ NiSOD
GSH
↑ GST
GSS-R
↑ TXN
$HCO_3^-$ + $NH_4^+$
unCPS (III)
carbamoyl-P
OTC
ornithine
citrulline
Urea Cycle
aspartate
ornithine
↑ AsuS
Arg
arginino succinate
arginine
AsL
AsL
fumarate

Figure S 6: **An overview of the proteomic response in the nitrogen and carbon metabolisms in *T. oceanica* (CCMP 1005) grown under Cu-limiting conditions.** Boxes indicate proteins with their abbreviated name. The colors of the boxes indicate expression in TO05 under low Cu: dark red, highly up-regulated (> 2-fold, $p<0.05$); light pink, up-regulated by 1.3 to 2-fold ($p<0.05$); dark blue, highly down-regulated (> 2-fold, $p<0.05$); light blue, down-regulated by 1.3 to 2-fold ($p<0.05$); white, expressed in TO05; grey border around box, found in *T. oceanica* genome but not detected in TO05 proteomic data; grey, dashed border around box, no putative homologs in the *T. oceanica* genome.

**Abbreviations:** AAT, aspartate aminotransferase; ACO, aconitase hydratase; Arg, Arginase; AsL, argininosuccinate lyase; AsuS, argininosuccinate synthase; cbbX, rubisco expression protein; CS, citrate synthase; DLDH, dihydrolipoamide dehydrogenase; EDA, 2-keto-3-deoxy phosphogluconate aldolase; EDD, 6-phosphogluconate dehydratase; ENO, Enolase; F2BP, Fructose-1-6-bisphosphatase; FBA I, fructose-bisphosphate aldolase class-I; FBA II, fructose-bisphosphate aldolase class-II; Fe-NiR, nitrite reductase (ferredoxin-dependent); FH, fumarate hydratase; GAPDH, glyceraldehyde 3-phosphate dehydrogenase; GDH, glutamate dehydrogenase; GOGAT, glutamate synthase; GPI, glucose-6-phosphate isomerase; GSII, glutamine synthetase; GSIII, glutamine synthetase; IDH, isocitrate dehydrogenase; MDH, malate dehydrogenase; NR, nitrate reductase; NRT, nitrate/nitrite transporter; OGD, 2-oxoglutarate dehydrogenase; OTC, ornithine carbamoyltransferase; PC, pyruvate carboxylase; PEPC, phosphoenolpyruvate carboxylase; PFK, phosphofructokinase; PGAM, phosphoglycerate mutase; PGK, phosphoglycerate kinase; PGM, phosphoglucomutase; PK, pyruvate kinase; RPE, ribulose-5-phosphate epimerase; RPI, ribose-5-Phosphate-isomerase; RuBisCO, ribulose-bisphosphate carboxylase; SUCLA, Succinate CoA synthetase; TPI, triose-phosphate isomerase; TPI, triose-phosphate isomerase; unCPS (CPSase III), carbamoyl-phosphate synthase; URT, Na/urea-polyamine transporter.

Table S 1: **Comparison of cellular localization of various carbon metabolic pathways.** (modified from (Gruber & Kroth, 2017)

| Pathway | Function | *Chlamydomonas* (green lineage) | Diatoms | Animals | Comment regarding diatoms |
|---|---|---|---|---|---|
| Calvin Cycle (Reductive Pentose P Way) | Carbon fixation 3RuBisP + 3 $CO_2$ -> GAP | Plastid | Plastid | -- | Rubsico has higher $CO_2$ affinity than other organisms |
| Oxidative Pentose P way | NAD(P)H, ribose (C5 sugars) for nt/ FA, AA | Plastid, Cytosol | Cytosol only (Kroth et al. 2008) | Cytosol | Cytosol only -> needed re-import of nts into plastid -> lots of nt transporters |
| Glycolysis 1$^{st}$ half (GAP) | Glucose -> GAP | Plastid | Plastid, Cytosol | Cytosol | |
| Glycolysis 2$^{nd}$ half (pyruvate) | GAP -> pyruvate | Cytosol (Johnson and Alric 2013) | Plastid, Cytosol, Mitochondria | Cytosol | Maybe also in mito to use Entner-Douderoff pathway generated pyruvate? |
| Entner-Douderoff pathway | 6PG ->pyruvate + GAP | -- | Mitochondria | -- | Fewer proteins involved (cost for cell), but also less ATP generated, but then again, faster as well, another tool to be adaptable to fast changing environment |
| Phosphoketolase pathway | Fructose-6-P -> acetyl-phosphate and GAP | -- | -- (only in *Phaeodactylum*) | -- | |
| Citrate Cycle | Carbon hub, reducing equivalents for respiration | Mitochondria | Mitochondria | Mitochondria | Integrated with urea cycle / aspartate shunt |
| Urea Cycle | Animals: catabolic nitrogen excretion Diatoms: anabolic nitrogen/carbon redistribution hub | -- | Mitochondria Cytosol | Mitochondria | Incorporation of recycled N and C; in combination with bacterial HGT proteins, production of polyamines involved in cell wall formation and the osmolyte proline |
| Malate Shunt | Redistribution of reducing equivalents and C4/5-compounds | Plastid, Mitochondria | Plastid, Mitochondria | Mitochondria | Often mentioned/proclaimed as part of cross-talk between chloroplast and mitochondria; no proteins found so far in Tp, only mitochondrial membrane proteins from oxaloacetate transporter in *P. tricornutum*. |
| Pyrimidine biosynthesis | | Plastid | Cytosol | Cytosol | They need to be transported across all four membranes |
| Carbon Storage | | Plastid, starch | Vacuole, Chrysolaminaran, lipids | Cytosol, lipid droplets (triglyceride) | |

Table S 2: **Differential expression and predicted cellular location of proteins involved in the Calvin-Benson-Bassham cycle** in TO03 and TO05 cultured in low Cu conditions *vs.* control.

| Calvin-Benson-Bassham Cycle | | | TO03[c] | | | | TO03[c] | | | |
|---|---|---|---|---|---|---|---|---|---|---|
| | | | original | | +EST | | original | | +EST | |
| Gene name[a] | Diatom homolog[b] | Name | sol | insol | sol | insol | sol | insol | sol | insol |
| cbbX, THAOC_24360 | Tp24123, Pt 36139 | cbbX, rubisco expression protein | **-2.25** | | **-2.25** | | -1.34 | -1.03 | -1.28 | -1.08 |
| rbcS | | rbcS, ribulose-bisphosphate carboxylase small chain | 1.12 | 1.23 | 1.14 | 1.2 | -1.63 | -1.49 | -1.62 | -1.44 |
| rbcL | | rbcL, ribulose-bisphosphate carboxylase large chain | 1.03 | 1.33 | 1.01 | 1.39 | -1.09 | -1.34 | -1.08 | -1.38 |
| THAOC_07617 | Tp270304 | PGK, phosphoglycerate kinase | 1.54 | -1.08 | | | **2.98** | -1.71 | | |
| THAOC_07617 contig_117075_1_571_+ | | | | | **6.85** | | | | 1.02 | |
| THAOC_07617 contig_55250_138_1396_+ | | | | | 1.89 | -1.16 | | | 1.17 | 1.05 |
| THAOC_13085 | Tp270231 | GAPDH, glyceraldehyde 3-phosphate dehydrogenase | -1.29 | **-4.16** | -1.43 | **-4** | 1.19 | -1.11 | 1.17 | -1.09 |
| THAOC_02438 | Tp30380, Pt54738, Fc274452 | TPI2, triose-phosphate isomerase | **3.3** | | **3.13** | | -1.06 | | -1.12 | |
| THAOC_35826 | Tp8590 | TPI1, triose-phosphate isomerase | 1.94 | | 1.91 | | 1 | | 1 | |
| THAOC_32006 | Tp8590 | TPI3, triose-phosphate isomerase | 1.45 | | 1.43 | | 1.09 | | 1.14 | |
| THAOC_00388 | Tp270396, PtBd825 | FBA II, fructose-bisphosphate aldolase class-II | 1.36 | | | | -1.23 | | | |
| THAOC_12069 | Tp21748, Pt22993 | FBA II, fructose-bisphosphate aldolase class-II | **2** | | 1.99 | | 1.15 | | 1.2 | |
| THAOC_02112 | no Tp, Pt51289 | FBA I, fructose-bisphosphate aldolase class-I | -1.73 | | -1.73 | | -1.18 | 1.08 | -1.35 | 1.18 |
| THAOC_31290 | | RPI, ribose-5-phosphate-isomerase | -1.12 | | -1.12 | | -1.09 | | -1.09 | |
| THAOC_31290 contig_19155_129_974_+ | | | | | **4.9** | | | | | |
| THAOC_09031 | | RPE, ribulose-5-phosphate epimerase | 1.22 | | -1.19 | | -1.1 | | -1.1 | |

[a]Gene name as per Lommer et al, 2012; if identified based on EST data, the respective contig identifier is given as well
[b]using blastp, this is the closest diatom homolog

[c]average fold-change in Cu-limited compared to control cultures, bold indicates highly differentially expressed (> ± 2-fold, $p<0.05$), underlined indicates differential expression ratio of ±1.3- to 2-fold ($p<0.05$). original, original proteomics dataset in which peptides where mapped to the TO05 genome; +EST, second proteomics dataset in which peptides where mapped to both the TO05 genome and our EST library; sol, soluble protein fraction; insol, insoluble protein fraction

Table S 3: **Proteins with triose-phosphate transporter PFAM and their expression in TO03 and TO05** in response to Cu limitation *vs.* control.

| Putative triose-P-transporters | | | TO03[c] | | | | TO05[c] | | | |
|---|---|---|---|---|---|---|---|---|---|---|
| | | | original | | +EST | | original | | +EST | |
| Gene name[a] | Diatom homolog[b] | TargetP/ASAFind | sol | insol | sol | insol | sol | insol | sol | insol |
| Cytosol | | | | | | | | | | |
| THAOC_31045 | triose-phosphate transporter | _(3) / Not plastid | | | | | | -1.08 | | -1.07 |
| THAOC_33538 | triose-phosphate transporter | _(2) / Not plastid | | | | | | | | |
| Chloroplast | | | | | | | | | | |
| THAOC_05244 | triose-phosphate transporter | S(2) / Plastid | | | | -1.82 | | -1.06 | | -1.06 |
| THAOC_11183 | triose-phosphate transporter | S(1) / Plastid | | | | | | | | |
| Mitochondria | | | | | | | | | | |
| THAOC_35890 | triose-phosphate transporter | M(3) / Not plastid | | | | | | 1.36 | | 1.36 |
| Signal Sequence | | | | | | | | | | |
| THAOC_04741 | triose-phosphate transporter | S(5) / Not plastid | | -1.31 | | -1.31 | | 1.34 | | 1.35 |
| THAOC_13820 | triose-phosphate transporter | S(3) / Not plastid | | | | | | | | |

[a]Gene name as per Lommer et al, 2012; if identified based on EST data, the respective contig identifier is given as well
[b]using blastp, this is the closest diatom homolog
[c]average fold-change in Cu-limited compared to control cultures, bold indicates highly differentially expressed (> ± 2-fold, $p<0.05$), underlined indicates differential expression ratio of ±1.3- to 2-fold ($p<0.05$). original, original proteomics dataset in which peptides where mapped to the TO05 genome; +EST, second proteomics dataset in which peptides where mapped to both the TO05 genome and our EST library; sol, soluble protein fraction; insol, insoluble protein fraction

Table S 4: **Differential expression and predicted cellular location of proteins involved in glycolysis** in TO03 and TO05 cultured in low Cu conditions *vs.* control.

| Glycolysis | | | TO03[c] | | | | TO05[c] | | | |
|---|---|---|---|---|---|---|---|---|---|---|
| | | | original | | +EST | | original | | +EST | |
| Gene name[a] | Diatom homolog[b] | Name | sol | insol | sol | insol | sol | insol | sol | insol |
| Cytosol | | | | | | | | | | |
| THAOC_06412 | Pt50444 | PGM, phosphoglucomutase | -1.79 | | **-3.17** | | -1.01 | 1.13 | -1.01 | 1.13 |
| THAOC_06412 contig_11427_139_3309_+ | | | | | -1.71 | | | | | |
| THAOC_25871 | Tp38266, Pt23924 | GPI, glucose-6-phosphate isomerase | -1.27 | | -1.26 | | 1.45 | | 1.34 | |
| THAOC_16559 | Tp22213 | PFK, phosphofructokinase | -1.8 | | -1.75 | | -1.03 | | 1.02 | |
| THAOC_05494 | Tp256250 | F2BP, fructose-1-6-bisphosphatase | 1.23 | | 1.21 | | -1.18 | | -1.21 | |
| THAOC_24977 | Tp270288, Pt29014 | FBA II, fructose-bisphosphate aldolase class-II | -1.18 | | -1.22 | | -1.32 | | -1.18 | |
| THAOC_24978 | Tp11761 | FBA I, fructose-bisphosphate aldolase class-I | -1.6 | | -1.53 | | -1.06 | **-2.37** | -1.09 | -2.26 |
| THAOC_24978 contig_128882_77_667_- | | | | | **-2.79** | | | | | |
| THAOC_21096 | Tp30380, Pt54738 | TPI2, triose-phosphate isomerase | -1.01 | | 1.04 | | 1.13 | | 1.25 | |
| THAOC_21095 | Pt54738 | TPI2, triose-phosphate isomerase | -1.25 | | -1.49 | | 1.01 | | -1.07 | |
| THAOC_34937 | Tp40393 | PK, pyruvate kinase | 1.58 | | 1.58 | | 1.28 | -1.61 | 1.25 | -1.61 |
| Chloroplast | | | | | | | | | | |
| THAOC_12069 | Tp21748, Pt22993 | FBA II, Fructose-bisphosphate aldolase class-II | **2** | | 1.99 | | 1.15 | | 1.2 | |
| THAOC_00388 | Tp270396, PtBd825 | FBA II, Fructose-bisphosphate aldolase class-II | 1.36 | | | | -1.23 | | | |
| THAOC_02112 | no Tp, Pt51289 | FBA I, Fructose-bisphosphate aldolase class-I | -1.73 | | -1.73 | | -1.18 | 1.08 | -1.35 | 1.18 |

| Glycolysis | | | TO03[c] | | | | TO05[c] | | | |
|---|---|---|---|---|---|---|---|---|---|---|
| | | | original | | +EST | | original | | +EST | |
| Gene name[a] | Diatom homolog[b] | Name | sol | insol | sol | insol | sol | insol | sol | insol |
| THAOC_02438 | Tp30380, Pt54738, Fc274452 | TPI2, triose-phosphate isomerase | **3.3** | | **3.13** | | -1.06 | | -1.12 | |
| THAOC_35826 | Tp8590 | TPI1, triose-phosphate isomerase | 1.94 | | 1.91 | | 1 | | 1 | |
| THAOC_32006 | Tp8590 | TPI3, triose-phosphate isomerase | 1.45 | | 1.43 | | 1.09 | | 1.14 | |
| THAOC_13085 | Tp270231 | GAPD1, glyceraldehyde 3-phosphate dehydrogenase | -1.29 | **-4.16** | -1.43 | **-4** | 1.19 | -1.11 | 1.17 | -1.09 |
| THAOC_07617 | Tp270304 | PGK, phosphoglycerate kinase | 1.54 | -1.08 | | | **2.98** | -1.71 | | |
| THAOC_07617 contig_117075_1_571_+ | | | | | **6.85** | | | | 1.02 | |
| THAOC_07617 contig_55250_138_1396_+ | | | | | 1.89 | -1.16 | | | 1.17 | 1.05 |
| THAOC_21902 | Tp28350 | PGAM2, phosphoglycerate mutase | **-2.25** | | **-2.25** | | -1.46 | | -1.19 | |
| Mitochondria | | | | | | | | | | |
| THAOC_33331 | Tp28334 | GAPDH, glyceraldehyde 3-phosphate dehydrogenase | **2** | -2.15 | 1.75 | | 1.42 | 1.19 | 1.29 | 1.32 |
| THAOC_33331 contig_105735_1_207_+ | | | | | **3.79** | | | | 1.55 | 1.3 |
| THAOC_33331 contig_105693_1_204_+ | | | | | | | | | 1.34 | |
| THAOC_07275 | Tp28241 | GAPDH, glyceraldehyde 3-phosphate dehydrogenase | 1.06 | | 1.04 | | 1.14 | 1.06 | 1.07 | 1.09 |
| THAOC_08963 | Tp27850 | PGAM, phosphoglycerate mutase | -1.03 | | -1.06 | | 1.12 | | 1.06 | |
| THAOC_34936 | Tp40391 | ENO, Enolase | 1.44 | | 1.54 | | 1.2 | 1.04 | 1.26 | 1.04 |
| THAOC_07097 | Pt561725 | PK, pyruvate kinase | -1.31 | | -1.31 | | -1.24 | -1.34 | -1.25 | -1.34 |
| THAOC_10009 | Pt34120, Fc267632, Tp38807 | EDA, 2-keto-3-deoxy phosphogluconate aldolase | 1.06 | | 1.06 | | 1.26 | | 1.26 | 1.32 |

[a]Gene name as per Lommer et al, 2012; if identified based on EST data, the respective contig identifier is given as well

[b]using blastp, this is the closest diatom homolog

[c]average fold-change in Cu-limited compared to control cultures, bold indicates highly differentially expressed (> ± 2-fold, $p<0.05$), underlined indicates differential expression ratio of ±1.3- to 2-fold ($p<0.05$). original, original proteomics dataset in which peptides

where mapped to the TO05 genome; +EST, second proteomics dataset in which peptides where mapped to both the TO05 genome and our EST library; sol, soluble protein fraction; insol, insoluble protein fraction

Table S 5: **Differential expression and predicted cellular location of proteins involved in the tricarboxylic acid (TCA) / citrate cycle** in TO03 and TO05 cultured in low Cu conditions *vs.*control. (for diagram, see Fig. 3, main text)

| <u>Citrate / TCA Cycle</u> | | | TO03[c] | | | | TO05[c] | | | |
|---|---|---|---|---|---|---|---|---|---|---|
| | | | original | | +EST | | original | | +EST | |
| NCBI protein identifier[a] | Diatom homolog[b] | Name | sol | insol | sol | insol | sol | insol | sol | insol |
| Mitochondria | | | | | | | | | | |
| THAOC_19912 | Tp11411, Pt30145 | CS, citrate synthase | 1.06 | | -1.05 | | <u>1.15</u> | | 1.07 | <u>1.14</u> |
| THAOC_20545 | Tp268965 | ACO, aconitase hydratase | **-4.73** | | **-4.17** | | <u>1.26</u> | | 1.3 | |
| THAOC_37807 | Tp21640, Pt14762, Fc168262 | IDH, isocitrate/ isopropylmalate dehydrogenase | <u>-1.64</u> | | <u>-1.66</u> | | <u>1.14</u> | | <u>1.14</u> | |
| THAOC_34595 | Tp1456, Pt45017 | IDH1, isocitrate dehydrogenase (monomeric) | **-3.02** | | **-2.79** | | <u>1.41</u> | | <u>1.45</u> | |
| THAOC_34595<br>contig_117709_1_1294_+ | | | | | **-5.5** | | <u>1.41</u> | | <u>-1.23</u> | |
| THAOC_28027 | Tp269718, Pt29016 | OGD1, 2-oxoglutarate dehydrogenase E1 component | 1.08 | | 1.08 | | <u>1.2</u> | | 1.03 | |
| THAOC_28027<br>contig_117377_84_1088_- | | | | | | | | | 1.34 | |
| THAOC_13425 | Fc226444 | SUCLA, succinate CoA synthetase, beta chain | 1.29 | | 1.29 | | <u>1.43</u> | | <u>1.57</u> | |
| THAOC_24310 | Tp24123, Pt 36139 | FH, fumarate hydratase, class II | -1.09 | | -1.09 | | <u>-1.23</u> | | <u>-1.18</u> | |
| THAOC_03405 | Tp20726, Pt51297 | MDH1, malate dehydrogenase | <u>1.6</u> | | <u>1.54</u> | | <u>1.15</u> | 1.05 | <u>1.15</u> | 1.05 |

[a]Gene name as per Lommer et al, 2012; if identified based on EST data, the respective contig identifier is given as well
[b]using blastp, this is the closest diatom homolog
[c]average fold-change in Cu-limited compared to control cultures, bold indicates highly differentially expressed (> ± 2-fold, $p$<0.05), underlined indicates differential expression ratio of ±1.3- to 2-fold ($p$<0.05). original, original proteomics dataset in which peptides

where mapped to the TO05 genome; +EST, second proteomics dataset in which peptides where mapped to both the TO05 genome and our EST library; sol, soluble protein fraction; insol, insoluble protein fraction

Table S 6: **Differential expression and predicted cellular location of proteins involved in nitrogen metabolism including the urea cycle** in TO03 and TO05 cultured in low Cu conditions *vs.* control.

| Nitrogen Metabolism | | | TO03[c] | | | | TO05[c] | | | |
|---|---|---|---|---|---|---|---|---|---|---|
| | | | original | | +EST | | original | | +EST | |
| Gene name[a] | Diatom homolog[b] | Name | sol | insol | sol | insol | sol | insol | sol | insol |
| Transporters | | | | | | | | | | |
| THAOC_04919 | Tp, Pt, Fc | NRT, nitrate/nitrite transporter | | **11** | | **11** | | | | |
| THAOC_01055 | Tp, Pt, Fc | NRT, nitrate/nitrite transporter | | | | | | <u>1.69</u> | | |
| THAOC_01056 | Tp, Pt, Fc | NRT, nitrate/nitrite transporter | | | | | | 1.36 | | <u>1.56</u> |
| THAOC_31656 | Tp24250, Pt20424, Fc214292 | URT, Na/urea-polyamine transporter | | **6.93** | | **6.93** | | 1.66 | | <u>1.32</u> |
| THAOC_07247 | Tp13996, Pt27877, Fc275907 | AMT, ammonium transporter | | -1.39 | | -1.39 | | -1.07 | | -1.11 |
| THAOC_00240 | Tp3393, Pt46427, Fc262069 | NiRT, formate/nitrite transporter | | <u>-1.14</u> | | <u>-1.14</u> | | <u>1.29</u> | | <u>1.14</u> |
| Cytosol | | | | | | | | | | |
| THAOC_34460 | Tp25299, Pt54983, Fc206583 | NR, nitrate reductase | <u>1.49</u> | | <u>1.45</u> | | -1.05 | <u>-1.32</u> | -1.05 | <u>-1.32</u> |
| THAOC_37170 | Tp42719, Pt, Fc207847 | AsuS, argininosuccinate synthase | 2.59 | | <u>-1.28</u> | | <u>1.39</u> | | <u>1.37</u> | |
| THAOC_12984 | Tp29075, Pt34526, Fc262171 | AsL, argininosuccinate lyase | -1.51 | | -1.51 | | 1.08 | | <u>1.12</u> | |

| Nitrogen Metabolism | | | TO03[c] | | | | TO05[c] | | | |
|---|---|---|---|---|---|---|---|---|---|---|
| | | | original | | +EST | | original | | +EST | |
| Gene name[a] | Diatom homolog[b] | Name | sol | insol | sol | insol | sol | insol | sol | insol |
| THAOC_04380 | Tp260953, Pt260953, Fc187231 | OCD, ornithine cyclodeaminase | 1.24 | | 1.24 | | 1.04 | | 1.22 | |
| THAOC_22587 | Tp34595, Pt25183, Fc190753 | ATCase, aspartate carbamoyltransferase | -1.05 | | -1.05 | | 1.2 | | 1.2 | |
| THAOC_06254 | Tp38359, Pt13951, Fc192073 | GDH, glutamate dehydrogenase, NADP dependent | 2.09 | | 2.77 | | 1.38 | | 1.5 | |
| THAOC_06254 contig_96116_1_202_- | | | | | 1.07 | | - | | 1.34 | |
| Chloroplast | | | | | | | | | | |
| THAOC_00016 | Tp262125, Pt27757, Fc185143 | NR, nitrite reductase | 1.34 | | 1.34 | | 1.07 | | 1.08 | |
| THAOC_02363 | Tp270365, Pt, Fc172194 | Fe-NiR, nitrite/sulfite reductase ferredoxin-like half-domain | **2.27** | | **2.32** | | -1.13 | | -1.1 | |
| THAOC_35252 | Tp26941, Pt, Fc200565 | NAD(P)H-NiR, nitrite reductase | -1.63 | | -1.65 | | 1.06 | | 1.07 | |
| THAOC_31900 | Tp26051, Pt51092, Fc228642 | GSII, glutamine synthetase | 1.71 | 1.76 | 1.59 | 1.81 | 1.27 | 1.46 | 1.28 | 1.5 |
| THAOC_31900 contig_127491_284_455_+ | | | | | **3.83** | | | | | |
| THAOC_13288 | Tp269900, Pt, Fc225787 | GOGAT, glutamate synthase | 1.55 | | 1.57 | | 1.05 | | 1.1 | |
| THAOC_16827 | Tp31394, Pt22909, Fc273803 | AAT, aspartate aminotransferase | **-2.33** | | **-2.33** | | 1.04 | | 1.03 | |
| THAOC_32466 | Tp270136, Pt50577, Fc277974 | argD, n-acetylornithine aminotransferase | -3.01 | | -2.97 | | 1.09 | | 1.09 | |
| Mitochondria | | | | | | | | | | |

Nitrogen Metabolism

| Gene name[a] | Diatom homolog[b] | Name | TO03[c] original sol | TO03[c] original insol | TO03[c] +EST sol | TO03[c] +EST insol | TO05[c] original sol | TO05[c] original insol | TO05[c] +EST sol | TO05[c] +EST insol |
|---|---|---|---|---|---|---|---|---|---|---|
| THAOC_17688 | Tp36208, Pt, Fc224733 | GDCT, glycine decarboxylase t-protein | -1.23 | | -1.23 | | | | | |
| THAOC_36273 | Tp39799, Pt22187, Fc276117 | GDCP, glycine decarboxylase p-protein | -1.86 | | -1.43 | | -1.17 | | -1.17 | |
| THAOC_01996 | Tp40323, Pt24195, Fc169332 | unCPS (CPSase III), carbamoyl-phosphate synthase | -1.14 | | -1.11 | | -1.06 | | -1.05 | |
| THAOC_05385 | Tp997, Pt30514, Fc268473 | OTC, ornithine carbamoyltransferase | 1.67 | | 1.67 | | -1.05 | | -1.05 | |
| THAOC_06032 | Tp270138, Pt22357, Fc277211 | GSIII, glutamine synthetase | **-5.29** | -2.62 | -7.01 | -2.62 | 1.34 | 1.16 | 1.18 | - |
| THAOC_06032 contig_17263_419_2602_- | | | | | | | - | | 1.37 | 1.22 |
| THAOC_15049 | Tp31424, Pt23059, Fc170472 | AAT, aspartate aminotransferase | **3.57** | | **3.57** | | | | - | |

[a]Gene name as per Lommer et al, 2012; if identified based on EST data, the respective contig identifier is given as well
[b]using blastp, this is the closest diatom homolog
[c]average fold-change in Cu-limited compared to control cultures, bold indicates highly differentially expressed (> ± 2-fold, $p$<0.05), underlined indicates differential expression ratio of ±1.3- to 2-fold ($p$<0.05). original, original proteomics dataset in which peptides where mapped to the TO05 genome; +EST, second proteomics dataset in which peptides where mapped to both the TO05 genome and our EST library; sol, soluble protein fraction; insol, insoluble protein fraction

Table S 7: **Differential expression and predicted cellular location of proteins involved in glutathione metabolism** in TO03 and TO05 cultured in low Cu conditions *vs.* control.

| Glutathione | | | TO03[c] | | | | TO05[c] | | | |
|---|---|---|---|---|---|---|---|---|---|---|
| | | | original | | +EST | | original | | +EST | |
| Gene name[a] | Diatom homolog[b] | Name | sol | insol | sol | insol | sol | insol | sol | insol |
| Cytosol | | | | | | | | | | |
| THAOC_29696 | Tp262753 , Pt47395 , Fc173923 | APX, ascorbate peroxidase | -1.17 | | -1.17 | | -1.27 | | -1.31 | |
| THAOC_37364 | Tp38724 , Pt___ , Fc___, Ehux444342 | APX, ascorbate peroxidase | 1.42 | | 1.37 | | 1.11 | 1.75 | 1.08 | 1.75 |
| THAOC_05931 | Tp267987 , Pt___ , Fc172515 | CYS2, cysteine synthase | -1.01 | | -1.01 | | 1.1 | | 1.06 | |
| THAOC_05931, contig_111591_1_205_- | | | | | | | | | 1.16 | |
| THAOC_05931, contig_68507_1_219_+ | | | | | -1.26 | | | | -1.02 | |
| THAOC_23355 | Tp13064 , Pt27240 , Fc291479 | GCS, γ-glutamyl-cysteine synthase | 4.53 | | 4.53 | | | | | |
| THAOC_09062 | Tp___ , Pt__ , Fc___, (*Hydra*, polyp) | GST, glutathione-S-transferase | **7.28** | | **12.4** | | | | | |
| THAOC_24526 | Tp269866 Pt37658 , Fc196320 | GST, glutathione-S-transferase | | | | | **2.11** | | **2.33** | 0.03 |
| THAOC_10112 | Tp40669 , Pt49037 , Fc267154 | NiSOD, nickel-dependent superoxide dismutase | 1.04 | 1.38 | 1.15 | 1.38 | 1.08 | 1.25 | 1.15 | 1.24 |
| THAOC_14269 | Tp___ , Pt49037 , Fc267154 | NiSOD, nickel-dependent superoxide dismutase | -2.68 | | -2.79 | | 1.54 | 1.15 | 1.54 | 1.15 |

| Glutathione | | | TO03[c] | | | | TO05[c] | | | |
|---|---|---|---|---|---|---|---|---|---|---|
| | | | original | | +EST | | original | | +EST | |
| Gene name[a] | Diatom homolog[b] | Name | sol | insol | sol | insol | sol | insol | sol | insol |
| THAOC_05213 | Tp41697 , Pt56471 , Fc231810 | TXN, thioredoxin | 2.62 | | **4.16** | | 1.33 | | 1.35 | |
| THAOC_06254 | Tp38359, Pt13951, Fc192073 | GDH, glutamate dehydrogenase, NADP dependent | 2.09 | | 2.77 | | 1.38 | | 1.5 | |
| THAOC_06254, contig_96116_1_202_- | | | | | 1.07 | | | | 1.34 | |
| Chloroplast | | | | | | | | | | |
| THAOC_27524 | Tp270338 , Pt_bd542 , Fc168185 | CYS, cysteine synthase | **2.43** | | **2.5** | | 1.38 | -1.44 | 1.28 | |
| THAOC_10442 | Tp267987 , Pt__ , Fc172515 | CYS2, cysteine synthase | 1.51 | | | | -1.3 | | -1.32 | |
| THAOC_10442, contig_37614_97_1300_+ | | | | | 1.61 | | | | 1.6 | |
| THAOC_19132 | Tp23356 , Pt36641 , Fc229440 | DHAR, dehydroascorbate reductase | | | | | | | | |
| THAOC_13288 | Tp269900 , Pt , Fc | GOGAT, glutamate synthase | 1.55 | | 1.57 | | 1.05 | | 1.1 | |
| THAOC_07268 | Tp26457 , Pt__ , Fc228584 | GR, glutathione reductase | **2.47** | | **2.4** | | 1.16 | | 1.34 | |
| THAOC_07269 | Tp11178 , Pt , Fc259160, Ehux58907 | GRX, glutaredoxin | -1.3 | | -1.3 | | 1.1 | | 1.1 | |
| THAOC_07269, contig_70896_7_411_- | | | | | -1.4 | | | | 1.01 | |
| THAOC_18234 | Tp270300 , Pt9035 , Fc225878 | GRX, glutaredoxin | -1.29 | | -1.4 | | 1.01 | | -1.24 | |

| Glutathione | | | TO03[c] | | | | TO05[c] | | | |
|---|---|---|---|---|---|---|---|---|---|---|
| | | | original | | +EST | | original | | +EST | |
| Gene name[a] | Diatom homolog[b] | Name | sol | insol | sol | insol | sol | insol | sol | insol |
| THAOC_04546 | Tp29212 , Pt25876 , Fc291481 | GSS, glutathione synthetase | 1.48 | | 1.48 | | -1.18 | | -1.27 | |
| THAOC_08875 | | GST, glutathione-S-transferase | | | | | 1.29 | | 1.29 | |
| THAOC_02860 | Tp32874 , Pt12583 , Fc 269011 | MnSOD, Mn/Fe binding superoxide dismutase | 1.77 | | 1.77 | | 1.07 | | 1.27 | |
| THAOC_31425 | Tp270230 , Pt46280 , Fc221387 | TXN, thioredoxin | 1.47 | | 1.47 | | 1.54 | | 1.64 | |
| Mitochondria | | | | | | | | | | |
| THAOC_02323 | Tpbd718 , Pt38124 , Fc182589 | GRX, glutaredoxin | 1.01 | | 1.01 | | 1.05 | | -1.11 | |
| THAOC_13865 | Tp19991 , Pt , Fc235520 | TXN, thioredoxin | 1.79 | | 1.77 | | 1.27 | | 1.26 | |

[a]Gene name as per Lommer et al, 2012; if identified based on EST data, the respective contig identifier is given as well

[b]using blastp, this is the closest diatom homolog

[c]average fold-change in Cu-limited compared to control cultures, bold indicates highly differentially expressed (> ± 2-fold, *p*<0.05), underlined indicates differential expression ratio of ±1.3- to 2-fold (*p*<0.05). original, original proteomics dataset in which peptides where mapped to the TO05 genome; +EST, second proteomics dataset in which peptides where mapped to both the TO05 genome and our EST library; sol, soluble protein fraction; insol, insoluble protein fraction

Table S 8: **Differential expression and predicted cellular location of proteins involved in the putative malate shunt** in TO03 and TO05 cultured in low Cu conditions *vs.* control.

| Malate-Shunt | | | TO03[c] | | | | TO05[c] | | | |
|---|---|---|---|---|---|---|---|---|---|---|
| | | | original | | +EST | | original | | +EST | |
| Gene name[a] | Diatom homolog[b] | Name | sol | insol | sol | insol | sol | insol | sol | insol |
| Transporter | | | | | | | | | | |
| THAOC_27515 | Tp26366, Pt23908, Fc180618 | malate-2-oxoglutarate antiporter | | | | | | 1.24 | | 1.24 |
| THAOC_25255 | Tp16746, Pt__, Fc226900 | malate-2-oxoglutarate antiporter | | | | | | 1.09 | | 1.09 |
| Mitochondria | | | | | | | | | | |
| THAOC_03405 | Tp20726, Pt51297 | MDH1, malate dehydrogenase | 1.6 | | 1.54 | | 1.15 | 1.05 | 1.15 | 1.05 |
| THAOC_19570 | no Tp homolog, Pt30519, Fc183259 | PC(1), pyruvate carboxylase | -1.63 | | -1.63 | | -1.14 | | -1.1 | |
| THAOC_15049 | Tp31424, Pt23059, Fc170472 | AAT, aspartate aminotransferase | **3.57** | | **3.57** | | | | | |
| THAOC_37124, contig_65096_132_669_+ | Tp34030, Pt56501, Fc277202 | ME, malic enzyme | | | | | | -1.15 | | |
| Chloroplast | | | | | | | | | | |
| THAOC_30817 | Tp25953, Fc291625 | MDH2, malate dehydrogenase, | **2.64** | | **2.64** | | | | | |
| THAOC_31413 | Tp269908, Pt49339, Fc186955 | PC(2), pyruvate carboxylase | **2.64** | | **2.7** | | 1.02 | | 1.01 | |
| THAOC_31413 contig_117322_1_422_- | | | | | 2.43 | | 1.02 | | 1.01 | |
| THAOC_16827 | Tp31394, Pt22909, Fc273803 | AAT, aspartate aminotransferase | **-2.33** | | **-2.33** | | 1.04 | | 1.03 | |

[a]Gene name as per Lommer et al, 2012; if identified based on EST data, the respective contig identifier is given as well
[b]using blastp, this is the closest diatom homolog

[c]average fold-change in Cu-limited compared to control cultures, bold indicates highly differentially expressed (> ± 2-fold, $p<0.05$), underlined indicates differential expression ratio of ±1.3- to 2-fold ($p<0.05$). original, original proteomics dataset in which peptides where mapped to the TO05 genome; +EST, second proteomics dataset in which peptides where mapped to both the TO05 genome and our EST library; sol, soluble protein fraction; insol, insoluble protein fraction

Table S 9: **Differential expression and predicted cellular location of proteins involved in respiration** in TO03 and TO05 cultured in low Cu conditions *vs.* control.

| Respiration | | | TO03[c] | | | | TO05[c] | | | |
|---|---|---|---|---|---|---|---|---|---|---|
| | | | original | | +EST | | original | | +EST | |
| Gene name[a] | Diatom homolog[b] | Name | sol | insol | sol | insol | sol | insol | sol | insol |
| THAOC_05862 | | NAD(P)H dehydrogenase (ubiquinone), alpha/beta subcomplex I | 1.91 | | 1.91 | | 1.21 | 1.42 | 1.47 | 1.48 |
| THAOC_04072 | | ubiquinol-cytochrome *c* reductase subunit VII | | -1.42 | | -1.34 | 0 | 1.24 | | 1.27 |
| THAOC_04852 | | cytochrome *c* oxidase subunit IV | | 1.52 | | 1.52 | 0 | **2.06** | | **2.06** |
| THAOC_22544 | | cytochrome *c* oxidase subunit Vb | -2.21 | 2.59 | -2.21 | 2.59 | -1.19 | **2.1** | -1.19 | 1.84 |
| THAOC_12811 | | cytochrome *c* oxidase subunit VIb | -1.15 | 4.03 | -1.15 | 4.03 | -1.12 | | -1.12 | |

[a]Gene name as per Lommer et al, 2012; if identified based on EST data, the respective contig identifier is given as well
[b]using blastp, this is the closest diatom homolog
[c]average fold-change in Cu-limited compared to control cultures, bold indicates highly differentially expressed (> ± 2-fold, $p<0.05$), underlined indicates differential expression ratio of ±1.3- to 2-fold ($p<0.05$). original, original proteomics dataset in which peptides where mapped to the TO05 genome; +EST, second proteomics dataset in which peptides where mapped to both the TO05 genome and our EST library; sol, soluble protein fraction; insol, insoluble protein fraction

**Notes S 1: Discussion on the contrasting adaptations to Cu limitation in the two strains of *T. oceanica* (CCMP1003 and CCMP1005).**

The response to Cu limitation is strikingly different between TO03 and TO05 (Figure S 5, Figure S 6). *Thalassiosira oceanica* TO03 is able to decrease its Cu requirement drastically by restructuring its complete photosynthetic apparatus (Hippmann et al, 2017) combined with complex changes in its carbon and nitrogen metabolism. These changes allow TO03 to alleviate ensued electron and ROS stress to sustain ongoing growth (this study). Strikingly, the main coping mechanism in TO05 to deal with Cu limitation lies in a 50% decrease of its entire proteome. Indeed, in response to low Cu, the only physiological parameters significantly affected in TO05 were growth rate, cell size and protein content (Hippmann et al, 2017).

In contrast to the response in TO03, none of the proteins involved in nitrogen acquisition and assimilation were highly up-regulated in TO05 (**Figure S 6**). The only proteins that were somewhat up-regulated (between 1.3- and 2-fold with $p<0.5$) were the involved transporters (URT, NRT, NiRT) and glutamine synthases (GSII/GSII). As neither NAD(P)H, $Fd^{red}$ oxidizing enzymes, or thioredoxins were up-regulated, it is unlikely that TO05 experienced any significant excess of reducing equivalents within the chloroplast, for which an enhanced nitrogen metabolism would be helpful. At this point, it is unclear how a potential increase in glutamine (both chloroplast and mitochondrial glutamine synthase are up-regulated) would be beneficial for the cell. It is also puzzling, why the Calvin-Benson-Bassham cycle influencing Rubisco activase, cbbX, and Rubisco itself were both somewhat down-regulated. This differs from the unchanged carbon assimilation rate in Cu-limited TO05. We acknowledge, though, that we have no information on metabolites in this study, so the hypothesis would be more robust if it were derived from combined results of significant changes in protein expressions and matching physiological data (e.g. inferred increased reduction state of chloroplast $Fe^{red}/Fed^{ox}$ in TO03 under Cu limitation).

The malate shunt was not induced under low Cu in TO05. Two homologs of a putative malate/oxoglutarate antiporter were expressed constitutively. However, chloroplast ATT and mitochondrial MDH1 were not even identified, suggesting levels below detection in at least one treatment (see methods). The two only indications of an active response to Cu limitation in TO05 is the significant up-regulation of glutathione-S-transferase and two glycolytic isoenzymes.

The differential expression of various isozymes involved in glycolysis is a central strategy in TO03 to direct energy and reducing equivalent flow within the cell (Section Carbon metabolism). TO05 does not exhibit the same level of sophistication in metabolic control. However, the only two significantly up-regulated proteins in carbon metabolism are chloroplast PGK and mitochondrial GAPDH, which would increase NADH in the mitochondria and ATP in the chloroplast. These two proteins might thus be the first line of defense in TO05 against small energy or reducing equivalent imbalances in the cell (**Error! Reference source not found.**).
